# Can single-component protein condensates form multiphase architectures?

**DOI:** 10.1101/2021.10.09.463670

**Authors:** Adiran Garaizar, Jorge R. Espinosa, Jerelle A. Joseph, Georg Krainer, Yi Shen, Tuomas P.J. Knowles, Rosana Collepardo-Guevara

## Abstract

Phase-separated biomolecular condensates that contain multiple coexisting phases are widespread in vitro and in cells. Multiphase condensates emerge readily within multi-component mixtures of biomolecules (e.g. proteins and nucleic acids) when the different components present sufficient physicochemical diversity (e.g. in inter-molecular forces, structure, and chemical composition) to sustain separate coexisting phases. Because such diversity is highly coupled to the solution conditions (e.g. temperature, pH, salt, composition), it can manifest itself immediately from the nucleation and growth stages of condensate formation, develop spontaneously due to external stimuli, or progressively as the condensates age. Here, we investigate thermodynamic factors that can explain the intrinsic transformation of single-component condensates into multiphase architectures during the nonequilibrium process of ageing. We develop a multiscale model that integrates atomistic simulations of proteins, sequence-dependent coarse-grained simulations of condensates, and a minimal model of dynamically ageing condensates with non-conservative inter-molecular forces. Our nonequilibrium simulations of condensate ageing predict that single-component condensates that are initially homogeneous and liquid-like can transform into gel-core/liquid-shell or liquid-core/gel-shell multiphase condensates as they age, due to gradual and irreversible enhancement of inter-protein interactions. The type of multiphase architecture is determined by the ageing mechanism, the molecular organization of the gel and liquid phases, and the chemical make up of the protein. Notably, we predict that inter-protein disorder-to-order transitions within the prion-like domains of intracellular proteins could lead to the required non-conservative enhancement of inter-molecular interactions. Our study, therefore, predicts a potential mechanism

**Significance Statement:** Biomolecular condensates are highly diverse systems spanning not only homogeneous liquid droplets, but also gels, glasses, and even multiphase architectures that contain various coexisting liquid-like and/or gel-like inner phases. Multiphase architectures form when the different biomolecular components in a multi-component condensate establish sufficiently imbalanced inter-molecular forces to sustain different coexisting phases. While such a requirement seems, at first glance, impossible to fulfil for a condensate formed exclusively of chemically-identical proteins (i.e., single-component), our simulations predict conditions under which this may be possible. During condensate ageing, a sufficiently large imbalance in inter-molecular interactions can emerge intrinsically from the accumulation of protein structural transitions—driving even single-component condensates into nonequilibrium liquid-core/gel-shell or gel-core/liquid-shell multiphase architectures.

## INTRODUCTION

Cells compartmentalize their interiors and regulate critical biological functions using both membrane-bound organelles and membraneless biomolecular condensates [1–3]. Condensates are ubiquitous mesoscopic assemblies of biomolecules that demix from the cytoplasm or nucleoplasm through liquid–liquid phase separation [1–5]. A dominant factor that enables biomolecules to phase separate is their multivalency: their ability to form multiple weak associative interactions [6–8]. Although not always present among phase-separating proteins[9], intrinsically disordered regions (IDRs)— often characterized by (amino acid sequences of low-complexity termed) low complexity domains, LCDs [10, 11]—have emerged as important contributors to the multivalency and phase separation capacity of many naturally occurring proteins [2, 3, 6]. When they have a high aromatic content, such as that found in prion-like domains (PLDs) [12, 13], IDRs can aggregate (fibrillation) and form amyloids [10, 11, 14]. Consistently, protein condensates that contain PLDs—for instance, the RNA-binding proteins Fused in Sarcoma (FUS), TAR DNA-binding protein 43 (TDP-43) and heterogeneous nuclear ribonucleoprotein A1 (hnRNPA1)—can undergo a further phase transition from functional liquid-like states to less dynamic reversible hydrogel structures or even irreversible gel-like states sustained by fibrillar aggregates [11, 15–23]. The transitions from liquid-like condensates to gel-like structures are implicated in neurodegenerative diseases such as amyotrophic lateral sclerosis and frontotemporal dementia [24–26].

Both within liquid-like and gel-like condensates, proteins interconnect forming percolated networks [27, 28]. However, gels can be distinguished from liquids by the prevalence of long-lived inter-protein interactions, which confer local rigidity to the network [27, 29]. The loss of the liquid-like character of a condensate over time, i.e. during ageing [15, 25], can be modulated by amino-acid sequence mutations, post-translational modifications, application of mechanical forces, and protein structural transitions, among many other factors [15–17, 20–22, 30–32]. These factors are expected to, directly or indirectly, increase the proportion of long-lived protein contacts within the condensate, which gives rise to complex mesoscale properties and network rigidity [27, 33].

Besides being highly diverse in terms of their material properties, condensates also vary significantly in their internal architectures [34]. Multi-component condensates can present various internal coexisting phases. The nucleolus [35], paraspeckles [35–37], and stress granules [38, 39], are all examples of hierarchically organized condensates with multiple coexisting phases or layers. Intranuclear droplets combining a dense liquid spherical shell of acetylated TDP-43—with decreased RNA-binding affinity—and an internal liquid core rich in HSP70 chaperones were recently observed [40]. Multicomponent multiphase condensates can also present internal low-density ‘bubbles’ or a ‘hollow’ space surrounded by an outer, denser phase [41–43]. Examples of these include the germ granules in Drosophila [41], the condensates formed from intracellular overexpression of TDP-43 [42], and in vitro RNA-protein vesicle-like condensates [43].

The multiphase behavior of condensates has been recapitulated in vitro, for instance in the liquid-core/gel-shell condensates formed by the Lge1 and Bre1 proteins [44], different multiphasic complex coacervates [45, 46], and multilayered RNA-protein systems [47, 48]. In all these cases, the emergence of a multilayered or multiphase organization is connected to the diversity in the properties of the biomolecular components within these condensates [28, 44, 46, 49]. Physicochemical diversity is key as it allows subsets of components to establish preferential interactions with one another leading to segregation into multiple layers inside the condensates, which are typically ordered according to their relative interfacial free energies [50]. Indeed, simulations and mean-field theory have shown that multi-component mixtures of species with sufficiently different valencies and/or binding affinities [51] are likely to segregate into multiple coexisting liquid phases with different compositions [52] or form multilayered architectures [46, 49, 50]. For instance, simulations of a minimal coarse-grained model recently showed that the diversity in the network of interactions in mixtures of unphosphorylated and phosphorylated FUS proteins can give rise to multiphase condensates [53]. In agreement with this idea, competing interactions among protein–RNA networks have been shown to drive the formation of multiphase condensates with complex material properties [51]. Experiments and simulations have further demonstrated that the different condensed liquid phases within multiphase condensates are hierarchically organized according to their relative surface tensions, critical parameters, viscosities, and densities [35, 46, 49, 50]. Motivated by these observations, here we explore the fundamental question of whether or not it is possible for single-component condensates to transition into a multiphase architecture as they age. We conceptualise condensate ageing as a nonequilibrium process where the inter-molecular forces among proteins exhibit gradual non-conservative changes over time.

Progressive dynamical arrest of single-component condensates has been observed in vitro and in cells for proteins with PLDs marked by low-complexity aromatic-rich kinked segments (or LARKS) [8, 10–20, 54, 55]. In these cases, formation of inter-protein β-sheets by LARKS peptides drives gradual fibrillation at the high protein concentrations present within condensates [18, 19, 23]. In such a situation, an imbalance in the inter-molecular forces among proteins inside the condensate is introduced and accumulates dynamically, resulting in nonequibrium behaviour. That is, rather than permanently establishing only weak, transient attractive interactions, some LARKS begin to assemble into inter-locking structures, which are strengthen due to the contribution of multiple hydrogen bonds and π–π interactions, leading to the formation of crossed-β-sheet amyloid structures [20]. Such a disorder-to-order transition within condensates highlights how subtle changes in the local behaviour of proteins can result in large-scale transformations of the mesoscale condensate structure.

Here, we investigate conditions that can dynamically alter the balance of inter-molecular forces among proteins within single-component homogeneous condensates, driving them out of equilibrium, and yielding multiphase architectures. To do so, we develop a multiscale modelling approach that allows us to connect multiple important scales of condensate formation, growth, equilibration, and nonequilibrium maturation. Our approach combines atomistic simulations of peptides with sequence-dependent coarse-grained simulations of protein condensates and a minimal model for protein condensates that age progressively. The minimal protein model is coupled to a dynamical nonequilibrium algorithm that mimics the progressive accumulation of inter-protein crossed-β-sheets during ageing. As a proof of concept, we focus on the naturally occurring phase-separating protein FUS, because it exhibits a liquid to gel transition during ageing[15, 30], and contains LARKS regions that can undergo a disorder-to-order transition to form β-sheet rich structures. Our simulations predict that FUS condensates can transition from equilibrium homogeneous condensed liquid phases into nonequilibrium liquid-core/gel-shell multiphase architectures due to the gradual enhancement in local protein interactions. Such enhancement can be provided, for instance, by an increase in cross-β-sheet content within LARKS regions of FUS proteins. Our simulations also demonstrate that the molecular organisation of the gel-like phase (e.g. whether the arrested state exposes more or less hydrophobic regions of the protein to the surface) significantly influences the relative interfacial free energy of the various protein phases; hence, determining the ordering of the various phases within multiphase condensates. Furthermore, the molecular organisation is also influenced by the ageing mechanism—e.g., which protein regions transition to form long-lived bonds dictates the preferential localization of residues that accumulate at the core or the surface of the gel phase. These findings highlight how variations in the binding strengths among proteins, due for instance to secondary structural changes, can result in nonequilibrium arrested multiphase condensates in single-component protein systems. These results further corroborate the idea that PLDs can act as modulators of prion-like protein phase behavior and, thereby, can tune collective interactions among adhesive amino acid motifs that result in condensate structural transformations that are observable at the mesoscale [12, 13].

## RESULTS AND DISCUSSION

### Multiscale model of single-component multiphase condensates

To provide mechanistic insights into the physical and molecular determinants that could drive single-component condensates to age into multiphase architectures, we designed a multiscale simulation approach that allows us to understand how subtle atomistic details of interacting proteins in solution impact the thermodynamic mechanisms of condensate formation and ageing (see Figure 1). Molecular simulations are powerful in dissecting the mechanisms, driving forces, and kinetics of phase separation and providing structural details of biomolecules within condensates, which are generally challenging to describe and interrogate using experimental techniques [56, 57]. All-atom and coarse-grained modelling approaches are now well-established tools, used in conjunction with experiments, to investigate biomolecular phase separation [28, 58–62]. Our multiscale modelling strategy leverages advantages and drawbacks of these two levels of modelling: (1) atomistic representations are used to describe the effects of chemical composition and protein structure in modulating inter-protein interactions, albeit in relatively small systems, and (2) coarse-grained models are developed and applied to consistently extrapolate such effects into the emergence of collective phenomena, such as condensate formation and ageing. Specifically, our approach integrates atomistic molecular dynamics (MD) simulations of interacting peptides (Step 1; Figure 1), amino-acid resolution coarsegrained simulations of protein condensates with implicit solvent (i.e., using our Mpipi protein model [63], which describes with near-quantitative accuracy the phase behaviour of proteins; Step 2; Figure 1), and a bespoke minimal coarse-grained model of protein condensates, in explicit solvent, that undergo ageing over time. To describe the nonequilibrium process of condensate ageing, we developed a dynamical algorithm that introduces dissipation through non-conservative inter-protein interactions. Specifically, the dynamical algorithm considers the gradual accumulation of stronger inter-protein bonds and local rigidification of the interacting protein segments due to inter-protein disorder-to-order transitions) inside the condensate (Steps 3–4; Figure 1). While our approach is fully transferable to other protein systems, as a proof of concept, we use it to investigate the ageing behaviour of single-component FUS condensates.

**FIG. 1:**
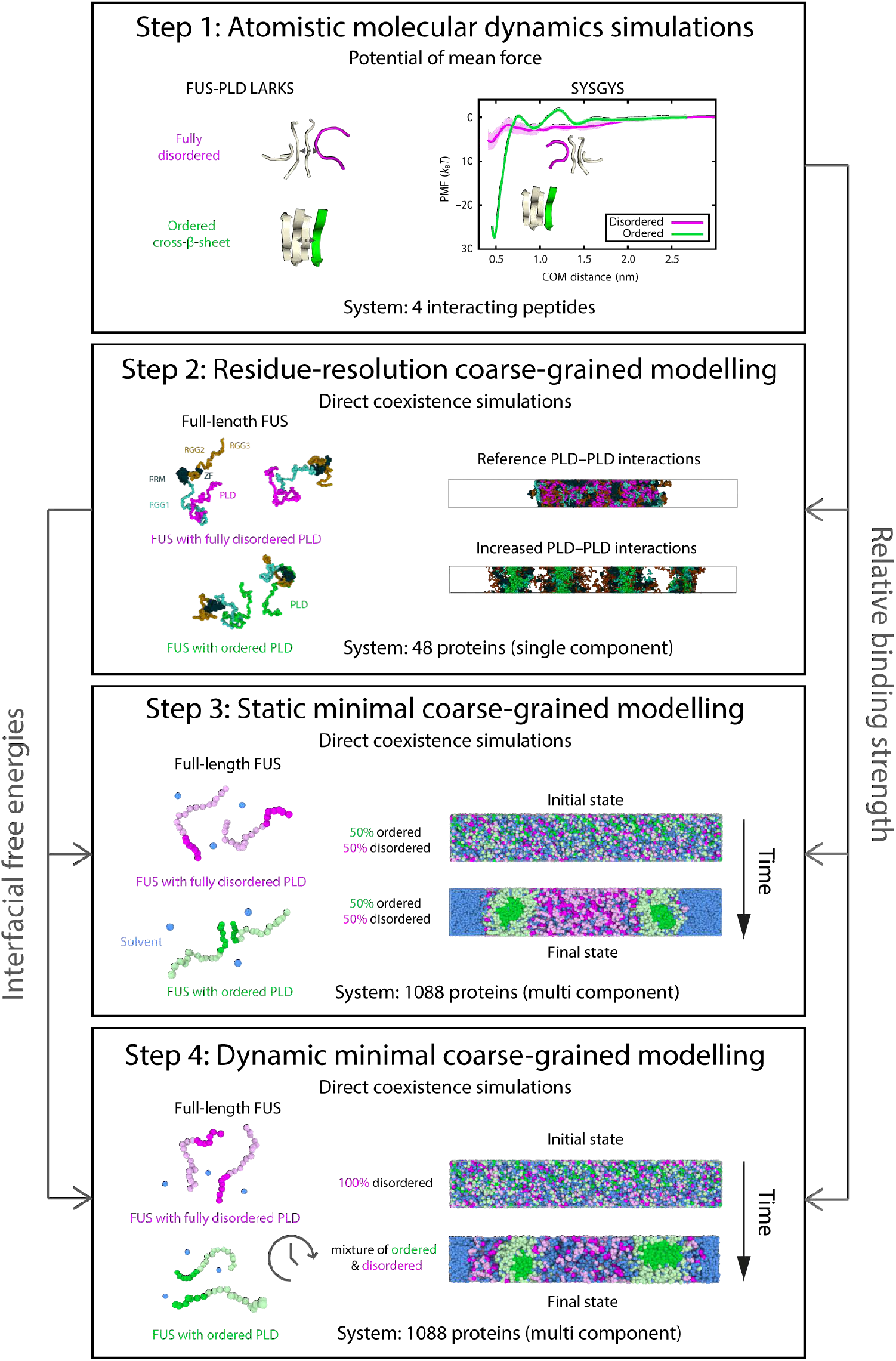
Visual summary of the multiscale computational approach employed in this work and divided into 4 steps explaining how the information from each step flows into the other steps. **Step 1**: Atomistic potential of mean force calculations (using Umbrella Sampling) are performed (for the different FUS short peptides that can undergo disorder-to-order transitions) to compare the free energy of binding when the sequences remain disordered *vs.* when they are structured. **Step 2**: With the information of the relative binding strength obtained in Step 1, the Mpipi coarse-grained model [63] is used to investigate the effect of increased PLD-PLD interactions in the interfacial free energy, molecular contacts and droplet organisation of FUS condensates. We find that strengthened PLD–PLD interactions drive the exposure of positively charged RGG2 and RGG3 domains to the droplet interface, thus, lowering their surface tension. **Step 3**: We develop a tailored minimal protein model to investigate FUS condensation in much larger system sizes (i.e., more than 20 times larger than with the Mpipi model). We design our tailored model based on the results from Steps 1 and 2, and observe the formation of steady-state multiphase FUS architectures. Finally in **Step 4**, we design a dynamic algorithm where instead of starting from a fixed concentration of structured *vs.* disordered FUS proteins, the disorder-to-order transitions spontaneously emerge over time depending on the protein concentration environment, and swapping the identity of proteins from being disordered to structured (with the same parameters as in Step 3) according to the PLD local coordination number. With this approach, nonequilibrium multiphase condensates are also observed consistent with findings of Step 3.

### A disorder-to-order structural transition diversify the interactions among chemically-identical FUS proteins

One of the proposed mechanisms used to explain ageing of RNA-binding proteins, like FUS, is their ability to undergo structural transitions within their LARKS. It has been observed that LARKS within FUS (e.g., SYSGYS, SYSSYGQS and STGGYG) can form pairs of kinked cross-β-sheets, which assemble into ladders and yield reversible fibrils that sustain FUS hydrogels [19]. Peptides at each step of the LARKS ladder form hydrogen bonds with adjacent peptides in the next step of the ladder. In addition, stacking of aromatic sidechains stabilize both the ladder and the individual β-sheets at each step, due to inter-molecular PLD–PLD interactions. We therefore hypothesized that the formation of cross-β-sheets among these critical LARKS in FUS-PLD could introduce sufficient physicochemical diversity into single-component FUS condensates to induce a multiphase organization.

To investigate this hypothesis, we first use atomistic MD simulations to compute the differences in interaction strength of FUS-PLD LARKS due to disorder-to-order transitions (Step 1; Figure 1). As done previously for Aβ-peptides [64], we perform atomistic umbrella sampling MD simulations to quantify the changes in the relative binding strengths (or Potential of Mean Force; PMF) among LARKS peptides when they are disordered *versus* when they stack to form the kinked β-sheet structures resolved crystallographically (Figure 2a) [19]. Following Ref. [65], we compute the inter-molecular binding strengths for three different FUS LARKS sequences (SYSGYS, SYSSYGQS, STGGYG) using two different force fields: a99SB-*disp* [66] (Figure 2) and CHARMM36m [67] (Figure S1). For each case, we calculate the free energy cost of dissociating a single peptide from a system containing four identical peptides. Four peptides is the smallest system size that considers the energetic cost of breaking both the step–step interactions and the peptide–peptide interactions within one step (i.e., there are two interacting peptides at each step of the ladder) upon single peptide dissociation.

**FIG. 2:**
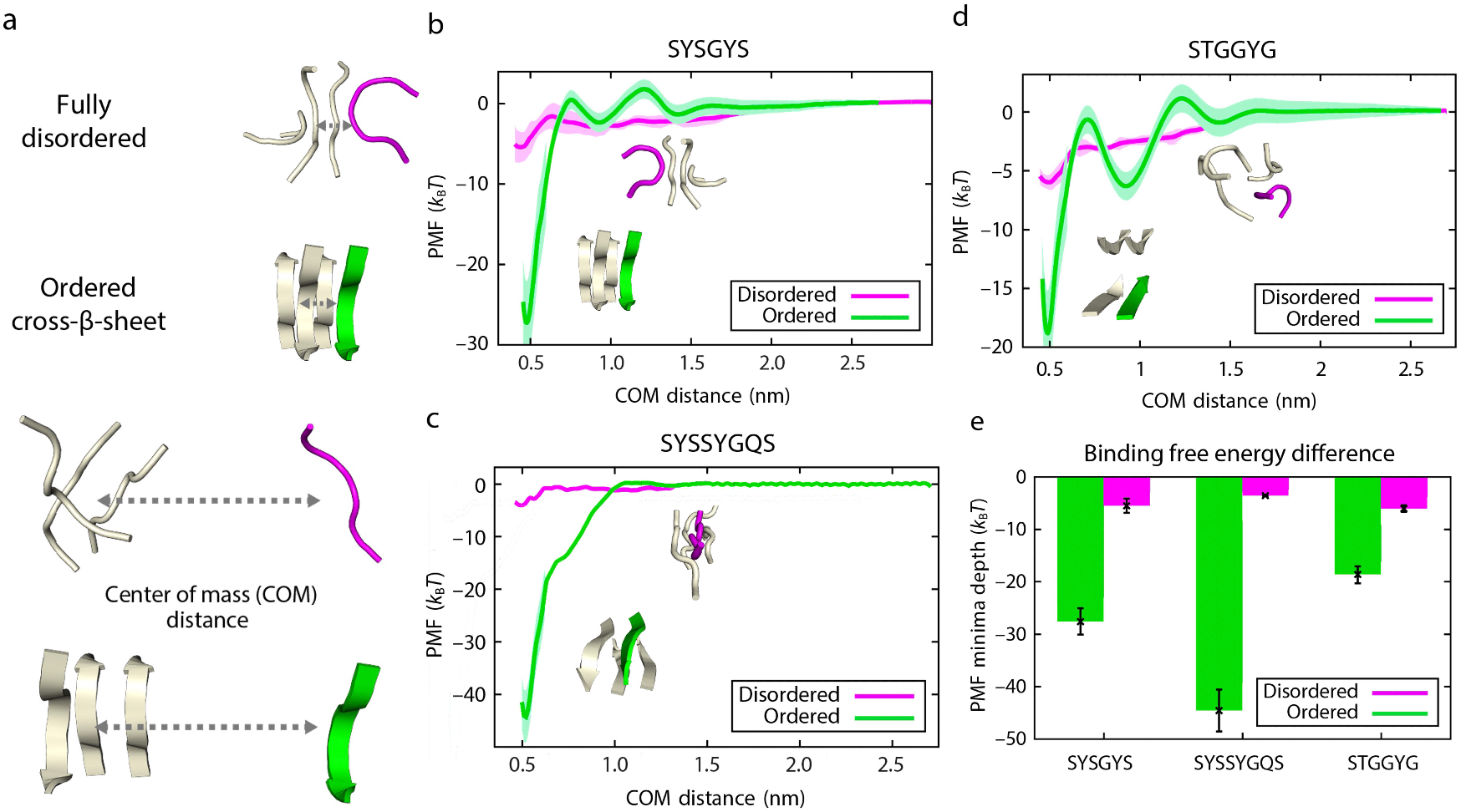
Disorder-to-order transition strengthens inter-molecular interactions among FUS LARKS. (a) Schematic of atomistic PMF simulations of LARKS-forming peptides in their disordered (magenta) and ordered states (green). The starting configurations for all simulations consisted of four stacked peptides. PMFs are calculated by estimating the free energy needed to remove one peptide (the ‘dissociating’ peptide) from the stack as a function of the centre of mass (COM) distance. The example shown is the peptide SYSGYS (PDB code: 6BWZ). (b–d) Plots of PMF *versus* COM for homotypic pairs of FUS LARKS-forming peptides SYSGYS (PDB code: 6BWZ), SYSSYGQS (PDB code: 6BXV) and STGGYG (PDB code: 6BZP), respectively, before (magenta) and after (green) undergoing the disorder-to-order structural transition. For the SYSGYS and STGGYG ordered systems, the PMFs display two barriers each (i.e., at COM distances of 0.7–0.8 and of 1.2–1.3 nm, respectively), which are contributed by the desolvation energy (primary and secondary shells) and the steric repulsion among the dissociating peptide and the remaining stacked peptides. In contrast, independently of the force field used (see Figure S1), the SYSSYGQS ordered system does not present noticeable steric repulsion among the dissociation path, and consistently, does not show energetic barriers at increasing values of the COM distance. Statistical errors, mean standard deviation, are shown as bands; obtained by bootstrapping the results from n = 5 independent simulations. (e) Variation in the free energy minimum (as obtained from the profiles in panel b–d)).

Independently of the force field used, the energetic cost of dissociating one of the peptides is ~four–eight times larger when these display canonical stacking, forming kinked β-sheet structures with two LARKS per step, *versus* when they are disordered. That is, when the four LARKS remain fully disordered, their binding interaction strengths is weak (~2–8 *k*_B_*T*), which suggests that thermal fluctuations can frequently break these interactions (Figure 2b–d and Figure S1). In contrast, when the LARKS peptides form β-sheets, the interaction strength increases by 15–40 *k*_B_*T* in total, depending on the LARKS sequence (Figure 2b–d and Figure S1). The increase is most noticeable in the SYSSYGQS system, likely due to the presence of Glutamine (Q); i.e., within the β-sheet structure Q exhibits an ideal orientation to act as an additional sticker by interacting strongly with both Serine (S) and Tyrosine (Y) [68]. A summary of the relative interaction strengths for the three LARKS sequences is provided in Figures 2e and S1d.

Overall our atomistic simulations suggest that the inter-molecular interactions among PLDs within a FUS condensate would be significantly strengthened upon formation of LARKS fibrillar-like ladders. However, in agreement with experiments, we find that the strength of interaction among such ordered LARKS is only sufficiently strong to sustain reversible hydrogels that dissolve upon salt treatment or heating, but not irreversible amyloids [19]. Irreversible amyloid fibrils would require larger binding energies, e.g. of the order of 50-80 *k*_B_*T* [19, 64, 69].

Based on the striking differences in the binding strength between FUS LARKS with β-sheets and those where the peptides remained fully disordered, we next investigated if a binary protein mixture composed of two distinct FUS conformational ensembles (i.e., with ordered *versus* with disordered LARKS) within the same condensate might give rise to a multiphase condensate morphology. In other words, we asked if two-phase coexistence within a single-component FUS condensate could emerge when a fraction of the FUS proteins have transitioned from having fully disordered PLDs to having PLDs that form inter-protein cross-β-sheets).

### Strengthening of PLD interactions dramatically transforms the molecular organization of FUS condensates

To determine the mesoscale implications of strengthening selected inter-protein bonds due, for instance, to an inter-molecular disorder-to-order transition, we perform amino-acid resolution coarse-grained simulations. In particular, we ask if such strengthening can give rise to FUS condensates with sufficiently different mesoscale properties from those of FUS condesates with standard interactions.

First, we use our Mpipi residue-resolution coarsegrained model [63] (Step 2; Figure 1), which recapitulates experimental phase separation propensities of FUS mutants [10], to characterize the molecular organization of single-component FUS condensates with standard (i.e., fully disordered and weakly interacting) PLD regions. Next, to probe the differences in the phase behavior of FUS proteins with PLDs that establish stronger interprotein interactions, we increase the PLD–PLD interactions as suggested by our atomistic PMFs, to approximate one of the key repercussion of inter-protein β-sheet formation. A representation of these residue-resolution coarse grained models for FUS are shown in Figure 3a, with the 526-residue FUS polypeptide chain sectioned into its PLD region (residues 1–165), three disordered RGG rich regions (RGG1: residues 166–267; RGG2: residues 371–421; and RGG3: residues 454–526), and two globular regions (an RNA-recognition motif (RRM): residues 282–371; and a zinc finger (ZF) domain: residues 422–453).

**FIG. 3:**
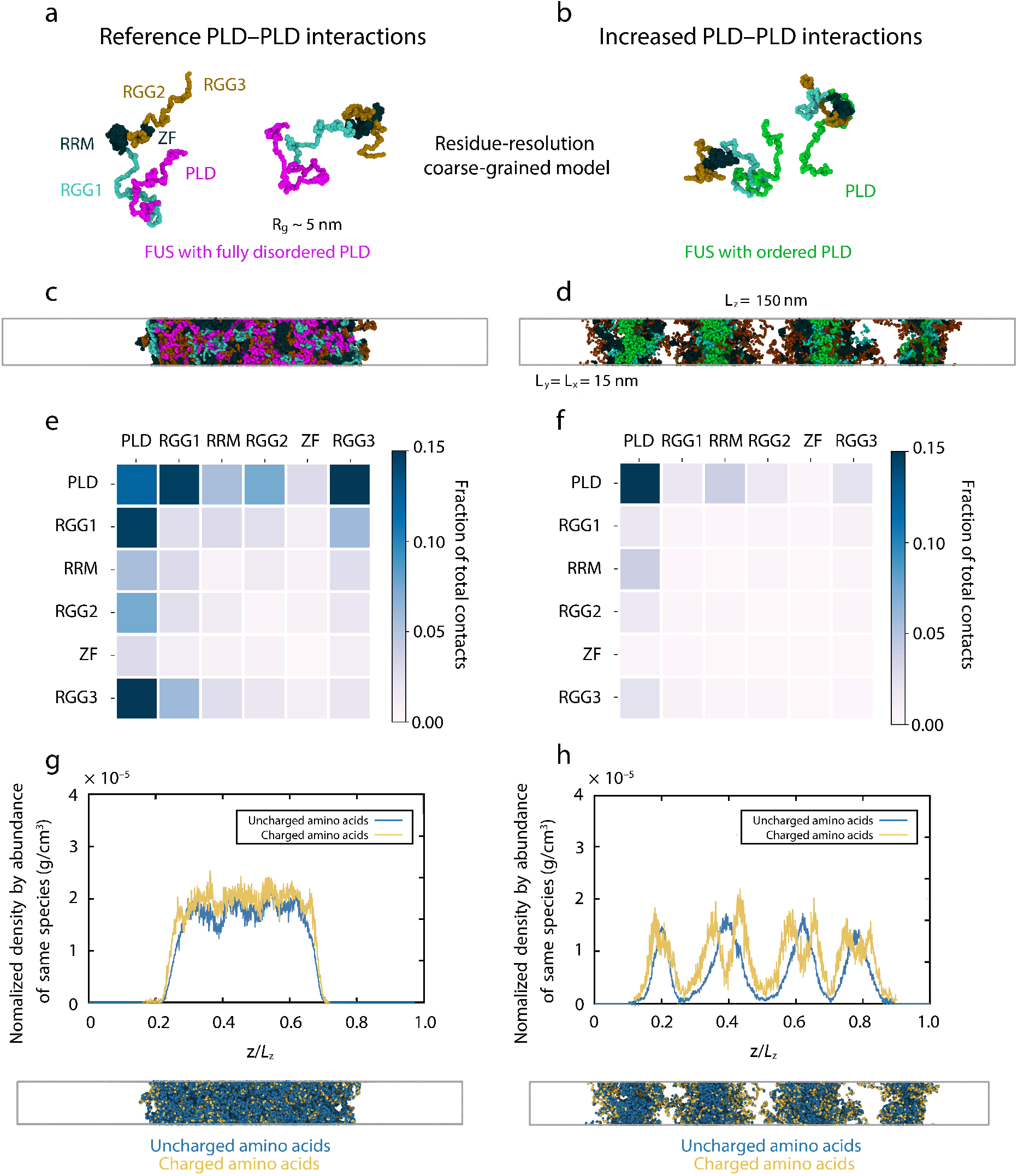
Strengthened PLD interactions give rise to multiphase condensate organization in coarse-grained molecular dynamics simulations. (a, b) Residue-resolution coarse-grained models for FUS with fully disordered PLD (panel a) and with ordered PLD (i.e., with cross-β-sheet elements in the PLDs) (panel b). Representative snapshots of FUS replicas, as obtained via direct coexistence MD simulations, are shown. Amino acid beads are colored according to the domains of FUS, with one bead representing each amino acid: PLD (residues 1–165): magenta (panel a) or green (panel b); RGG1 (residues 166–284): cyan; RGG2 (residues 372–422) and RGG3 (residues 454–526): ochre, RRM (residues 285–371) and ZF (residues 423–453): dark blue. The single-protein radius of gyration (*R_g_*) of FUS within the dilute phase (and therefore with reference PLD-PLD interactions) at the same conditions of the rest of simulations (*T* ~ 0.9*T_c_*) is also included. (c, d) Snapshots of direct coexistence simulations with reference interaction strengths among PLDs (panel c) and increased interactions strengths among PLDs (panel d) The simulation box sides included in panel d also apply for panels c, g and h. 48 FUS proteins were included in the simulations. Color code as in panel a and b. (e, f) Frequency of contacts between FUS domains within condensates for simulations with standard interaction strengths among PLDs (panel e) and increased interactions strengths among PLDs (panel f). Heatmaps are color coded and scaled from white to dark blue. The statistical uncertainty in the fraction of molecular contacts is 0.01. (g, h) Normalized density of charged (yellow) and uncharged (blue) species across the long side of the simulation box estimated over the coarse-grained equilibrium ensemble for simulations with standard interaction strengths among PLDs (panel c) and increased interactions strengths among PLDs (panel d). Density uncertainty corresponds to a 5%. The snapshots from direct coexistence simulations (bottom) are the same as in panel c and d, but now colored-coded according to the charge state of the amino acid residues.

For each of these coarse-grained parameterizations, we conduct residue-resolution direct coexistence simulations [70–72] of tens of interacting full-length FUS molecules, and estimate the influence of modulating the inter-molecular interactions among PLDs on FUS phase separation (Step 2; Figure 1). The direct coexistence method enables simulating a protein-enriched condensed liquid phase in contact with a protein-poor diluted-liquid phase in one simulation box and, thus, can determine whether a system phase separates at a given set of conditions.

Consistent with previous observations [73], FUS condensates formed by standard proteins with fully disordered PLDs display a homogeneous molecular architecture (Figure 3c,g) (i.e., all FUS domains are randomly positioned throughout the condensate). In contrast, FUS condensates containing strengthened PLD–PLD interactions (Figure 3d,h) exhibit a micelle-like heterogeneous organization with a PLD-rich hydrophobic core and a charge-rich interface; i.e., the positively charged RGG2 and RGG3 regions are effectively exposed to the solvent. As a result, the surface tension (*γ*) of FUS condensates with strengthened PLD–PLD interactions (*γ* = 0.06 ± 0.04*mJ/m*^2^) is considerably lower than that of the condensates formed by standard FUS proteins (*γ* = 0.32 ± 0.06*mJ/m*^2^). Consistently, FUS condensates with strengthen PLD–PLD interactions can stabilize multiple droplets in the same simulation box (Figure 3d,h), as observed previously for condensates with a surfactant-rich interface [50].

### Presence of distinct FUS structural ensembles supports the formation of hollow liquid-core/gel-shell condensates

To investigate the phase behaviour of a multicomponent condensate that contains both FUS proteins with fully disordered PLDs (termed ‘disordered’ FUS herein) and FUS proteins where the PLDs form inter-protein cross-β-sheets (termed ‘ordered’ FUS herein), we developed a minimal model (Step 3; Figure 1). A minimal model significantly reduces the degrees of freedom of the system, and hence the computational cost, while retaining essential physicochemical information. Such a reduction is required because simulations of multicomponent condensates must consider a larger number of proteins (~10^3^) than those of single-component systems to address the additional finite-size effects associated with approximating the overall composition of the system (i.e. the relative number of copies of each component).

In this new minimal model, full-length FUS proteins are each represented as a chain of 20 beads (i.e., 6 beads for FUS-PLD, and 14 beads for the RGG1, RRM, RGG2, ZF, and RGG3 regions). To distinguish between disordered and ordered FUS proteins, our minimal model considers the following three key physicochemical differences between the ordered and disordered FUS proteins, based on the results from our atomistic and residue-resolution simulations: (1) While disordered FUS proteins are modelled as fully flexible chains, we increased the local rigidity among PLD beads within ordered FUS proteins to mimic the structural effect of cross-β sheet formation. (2) We parameterize the strengths of interactions among pairs of ordered and disordered FUS domains differently. In each case, we set the strength of interactions among individual FUS regions (e.g., PLD– PLD, PLD–RGG1, RGG1–RGG1, etc.) using the relative frequencies of interactions obtained in their respective residue-resolution simulations; i.e., the disordered FUS minimal model according to the contact maps of the single-component standard FUS simulations (Figure 4a), and the ordered FUS model based on the contact maps of our single-component FUS simulations where the PLD–PLD interactions were increased based on our atomistic results (Figure 4b). (3) To recover the lower surface tension we observed in our residue-resolution simulations for single component ordered FUS condensates, we assign a higher hydrophilicity to the charged-rich FUS regions (i.e., RGG1, RGG2 and RGG3 regions) than to the other FUS domains, and incorporate a compatible minimal explicit solvent model. As a test, we run two separate simulations of single-component FUS condensates, one using the minimal model for ordered FUS and the other using the minimal model for disordered FUS (Figure S7). This test reveals that the minimal model is able to recapitulate both the micelle-like (for ordered FUS) and homogeneous (for disordered FUS) condensate organisations that we predicted with the residue-resolution simulations using similar system sizes and simulation conditions (Figure S7).

**FIG. 4:**
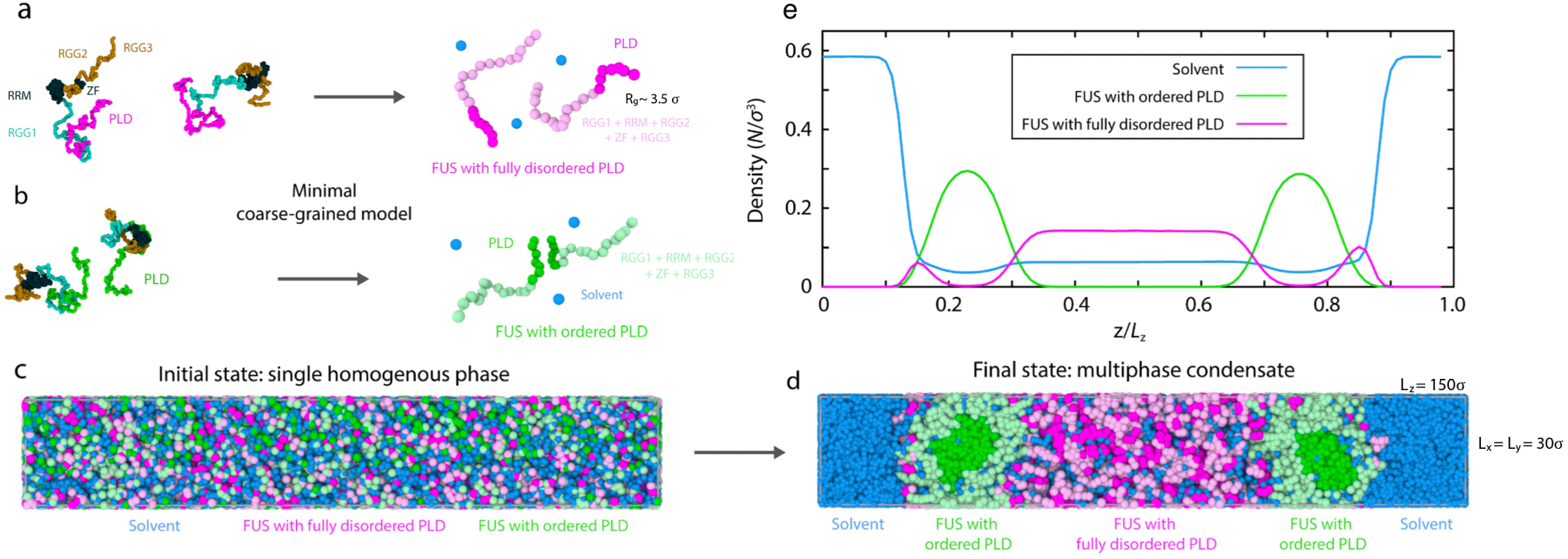
A disorder-to-order transition within PLDs drives multiphase organization in FUS condensates. (a, b) Minimal coarse-grained models for FUS with fully disordered PLDs (panel a) and with ordered PLDs (i.e., with cross-β-sheet elements in the PLDs) (panel b). Representative snapshots of FUS replicas, as obtained via direct coexistence MD simulations, are shown. Here one bead represents 26 amino acids. PLD: magenta (panel a) or green (panel b), RGG1, RRM, RGG2, ZF, RGG3: light magenta (panel a) or light green (panel b). Solvent (water) is depicted by blue beads. The radius of gyration (*R_g_*) of the minimal FUS coarse-grained protein in the dilute phase is ~3.5 *σ* (see information about reduced units in the Supporting Information (SI) Appendix). (c, d) Snapshots of direct coexistence simulations of a mixture of 50% FUS proteins with fully disordered PLDs and 50% FUS proteins with cross-β-sheet elements in their PLDs and explicit solvent. The initial state is shown in panel c and the final state in panel d. 1088 FUS proteins were included in the simulations. (e) Density profile (in reduced units) of FUS species and explicit solvent across the long side of the simulation box estimated over the coarse-grained equilibrium ensemble (as obtained in panel d). FUS proteins with fully disordered PLDs: magenta; FUS proteins with ordered PLDs (i.e., with kinked cross-β-sheets): green, solvent (water): blue.

Using these two minimal FUS protein models, we now perform direct coexistence simulations of a mixture of 50 mol % ‘disordered’ FUS proteins and 50 mol% ‘ordered’ FUS proteins in explicit water/solvent. By starting from a homogeneous mixed phase (Figure 4c), the system readily demixed into a phase-separated condensate with an inhomogeneous organization (Figure 4d). Evaluation of the density profile of the steady-state system reveals the formation of a liquid-core/gel-shell (i.e., hollow) multiphase condensate architecture; i.e., condensates are hierarchically organized with a low-density core phase, made up by ‘disordered’ FUS proteins, which is surrounded by a high-density shell composed of ‘ordered’ FUS species (Figure 4e). The lower density of the inner FUS phase with disordered PLDs is evident from its higher water content. Our simulations further reveal that FUS proteins in the outer shell are less mobile than in the inner core, as gauged by their reduced mean squared displacement (Figure S3). We verified that variations in the stoichiometry of the disordered FUS proteins *versus* ordered FUS proteins (Figure S5), as well as in the simulation system size and box shape (Figure S8), barely affect the organization of the liquid-core/gel-shell multiphase condensates. Together, our simulations predict that a FUS condensate that contains both a population of proteins that remain fully disordered and bind weakly, and of proteins that establish strong PLD–PLD interactions (i.e. due to the ordering and stacking into kinked cross-β-sheet fibrillar-like LARKS ladders) can self-assemble into a multiphase architecture.

### Ageing simulations predict multiphase condensate formation from single-component systems

The highly concentrated environments of biomolecular condensates may serve as seeding grounds for disorder-to-order transitions that occur in FUS-PLD LARKS (Step 4; Figure 1), or other mechanisms that produce local enhancement of inter-protein interactions. To gain deeper insights into the mechanism of FUS multiphase condensate formation and ageing, we next developed a dynamical algorithm that approximates the effects (i.e., strengthening of inter-protein bonds, local protein rigidification, and changes in the molecular organization of the condensed phase) of the gradual accumulation of interprotein β-sheet structures in a time-dependent manner and as a function of the local protein density. Rather than imposing a fixed concentration of ‘disordered’ and ‘ordered’ FUS proteins in the simulation a priori (as done in the previous section), this algorithm enabled us to study the spontaneous emergence of multiphase FUS condensates during ageing.

We again perform direct-coexistence simulations using our minimal FUS protein model, but now starting from a system containing 100% FUS with disordered PLDs and explicit solvent. As time progresses, our dynamical algorithm triggers disorder-to-order transitions within the PLD of FUS when high local fluctuations of protein densities are found within the condensate (Figure 5a). Specifically, transitions of LARKS into kinked cross-β-sheets are enforced and recapitulated by modulating the binding interaction strength of PLD–PLD interactions by a factor of eight (according to our atomistic PMF simulations) when a PLD is in close contact with four other PLDs and still possesses enough free volume around it to undergo a disorder-to-order structural transition (i.e., solvent-mediated) [74, 75] (see SI Appendix for justification and details). An important feature of this dynamic algorithm is that it allows us to explore ageing as a nonequilibrium process where the strength of interprotein interactions in the system is non-conservative (i.e. transitions from the strongly-interacting ordered-LARKS state back to the weakly-binding disordered state are for-bidden).

**FIG. 5:**
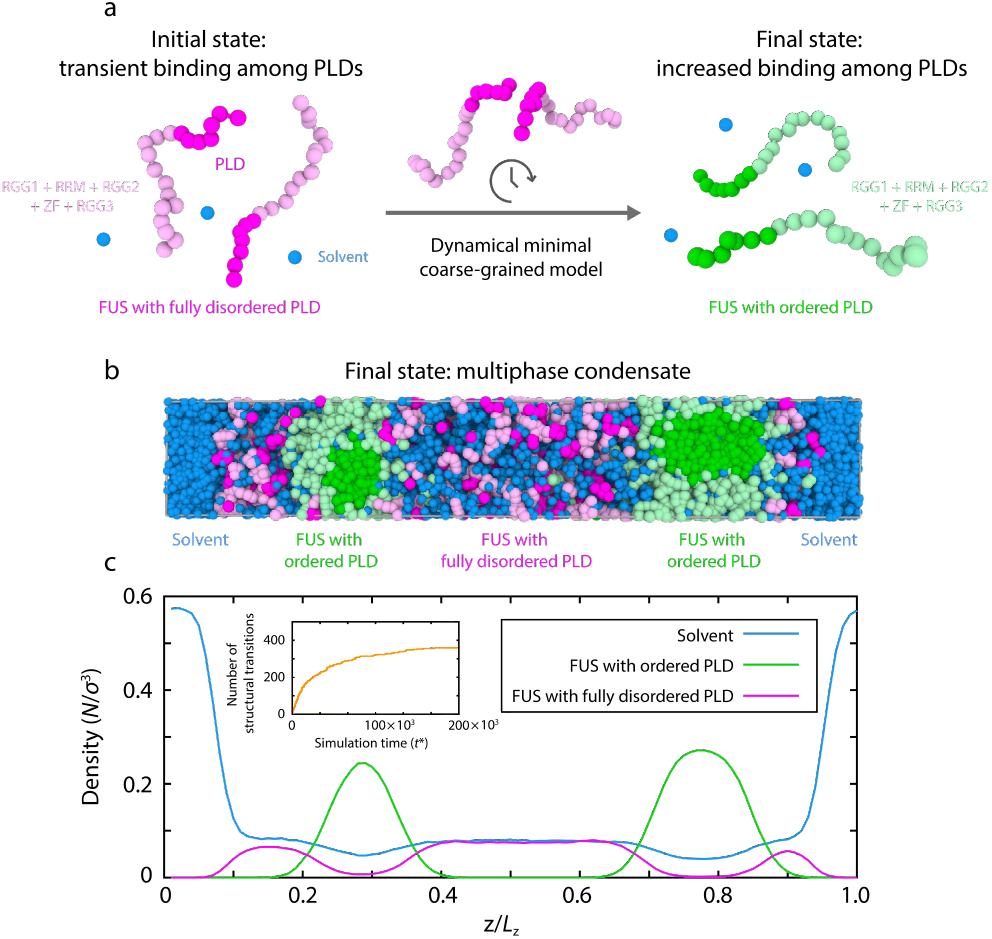
An ageing model predicts multiphase FUS condensates. (a) Schematic illustration of the dynamical algorithm for triggering disorder-to-cross-β-sheet transitions in the minimal coarse-grained model of FUS. This algorithm fosters disorder-to-order transitions by increasing the PLD–PLD interaction strengths by a factor of 8 (according to our atomistic PMF simulations) when FUS-PLDs are in close contact with other four other PLD-FUS domains and still possess enough surrounding free volume to undergo the transition into kinked cross-β-sheets. PLD: Magenta or green; RGG1, RRM, RGG2, ZF, RGG3: light magenta or light green. Solvent (water) is depicted as blue beads. (b) Snapshot of direct coexistence simulations using the dynamical algorithm at a quasi-equilibrium state after structural transitions have saturated. (c) Density profile (in reduced units) of FUS species and explicit solvent across the long side of the simulation box estimated over the coarse-grained quasi-stationary state (as obtained in panel b). Inset: Number of structural transitions in FUS-PLD domains as a function of simulation time (*t*^∗^). The plateau region was used to calculate the snapshots and density profiles in panel b and c, respectively. FUS proteins with fully disordered PLDs: magenta; FUS proteins with ordered PLDs (i.e., with kinked cross-β-sheet elements): green; solvent (water): blue.

Triggering the dynamic algorithm drives the initially homogeneous condensate to adopt a nonequilibrium multiphase architecture with a high-density outer shell and a low-density inner core (Figure 5b–c). The resulting multiphase architecture is equivalent to the one obtained in the previous section, when combining ordered and disordered FUS proteins at fixed concentrations (c.f., Figure 4). Comparing the phase diagram (in the plane of reduced temperature *versus* density) of a standard FUS condensate (i.e., with fully disordered FUS proteins; static algorithm) with that of a FUS condensate that has aged progressively over time (i.e., simulated with our dynamical algorithm) reveals only a moderate increase in the global density of the condensate upon ageing, but almost no change in the maximum temperature at which phase-separation is observed (Fig. S6).

Regarding the ageing mechanism, our dynamic algorithm predicts an initial steep increase in the rate of disorder-to-order transitions for FUS inside the condensate due to the high probability of local high-density fluctuations (Figure 5c inset and phase diagram in Fig. S6). Once the multiphase condensate begins to form, the rate of emergence of structural transitions decays rapidly, until a quasi-dynamically arrested state is reached. The dominant factor slowing down the rate of structural transitions comes from the formation of the low-density inner core; i.e., the lower densities in such a region significantly decrease the probability that clusters of four or more peptides would form. As the multiphase architecture consolidates even further, a slight speed up to the rate of transitions comes from the increasing numbers of ordered FUS proteins that become available at the newly formed shell–core interphase. These exposed ordered proteins can target disordered proteins from the low-density phase and drive them to undergo disorder-to-order transitions. However, the latter speed up is frustrated by the slower dynamics of the ordered FUS proteins, and by the steric barrier that FUS domains adjacent to the ordered PLD regions pose; or alternatively when significant modifications to the parameter set of the dynamic algorithm controlling the emergence rate of structural transitions are applied (Figure S4).

### Formation of liquid-core/gel-shell versus gel-core/liquid shell condensates

While FUS forms hollow condensates (Fig. 5b)— with a liquid-core/gel-shell architecture—we reasoned that other proteins may form different steady-state multiphase architectures combining gels and liquids. In other words, the specific ordering of the gel phase as the outer layer in FUS emerges from its amino acid sequence and the molecular organization of the various FUS domains inside its gel phase—i.e., as strong LARKS–LARKS bonds form, the charged-rich RGG regions of FUS are pushed towards the surface of the gel, making the gel more hydrophilic than the inner liquid core.

To investigate how the liquid and gel phases may organize in condensates formed by other proteins beyond FUS, or in FUS undergoing a different ageing mechanism, we let the surface of the gel to be equally (or less) hydrophilic than that of the liquid-like condensed phase. For this, we devise a set of control simulations where we now define all domains within our minimal protein model as equally hydrophilic. Using this model (see parameterization in Table 1 and 2 of the Methods Section), we perform two types of direct coexistence simulations: (1) for a system where we mix a priori 50% of fully ‘disordered’ proteins with 50% ‘ordered’ proteins with structured LARKS, and (2) for a system that is formed initially by 100 mol% ‘disordered’ proteins and where a protein region can dynamically undergo disorder-to-order transitions (as in Figure 5). Importantly, these new simulations predict that the condensates will now exhibit a nonequilibrium multiphase gel-core/liquid-shell arquitecture (Figure 6a); i.e., where the gel-like aged phase is now preferentially located in the core and the liquid forms an outer shell around it. These simulations suggest that the specific ordering of the gel and liquid phases is determined by the relative hydrophilicity of the coexisting gel and liquid phase surfaces. Given equal hydrophilicity of both gel and liquid surfaces, the gel, unsurprisingly, accumulates at the core due to the stronger protein–protein interactions that sustain it, and the lower surface tension of the liquid-like phase with the surrounding solvent [50].

**FIG. 6:**
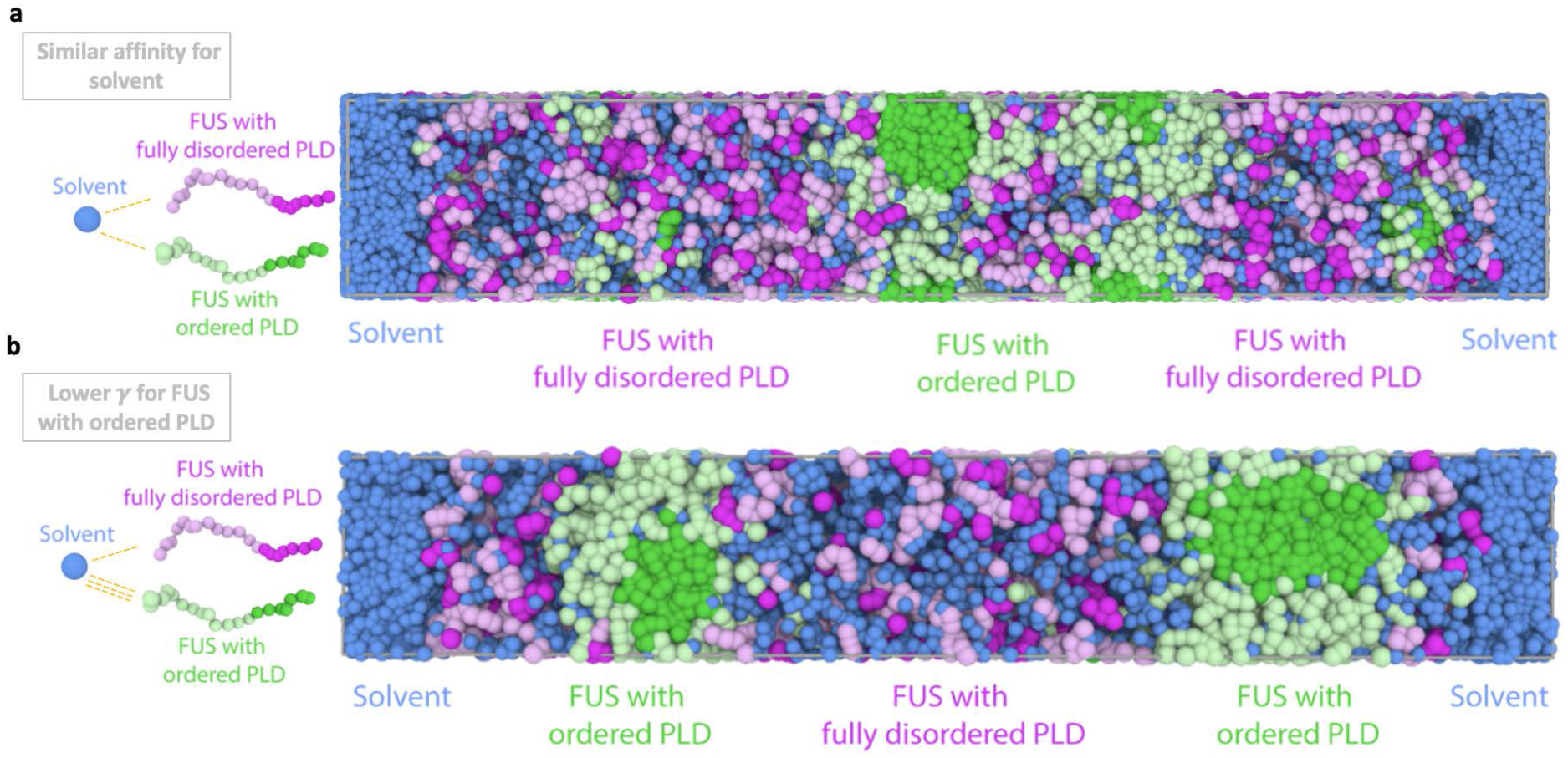
Equal hydrophilicity for structured and disordered PLD FUS proteins causes gel-like phase to relocate to the core of multi-phase condensates. Final state comparison between a control simulation with the dynamical algorithm (a) where we set the same solvent–protein interactions regardless of protein type (for both ordered and disordered PLDs) and (b) the dynamical algorithm simulations from Figure 5 where FUS with ordered PLDs present lower surface tension due to the exposure of their RGG regions to the interface of the droplets. Left: Schematic illustration of solvent–protein interactions. Right: (a) Snapshot of the quasi-equilibrium state of a Direct Coexistence simulation where fully disordered FUS and FUS with ordered PLDs exhibit equal hydrophilicity (i.e. they have the same affinity for water.) (b) Snapshot of the quasi-equilibrium state of a Direct Coexistence simulation where FUS with ordered PLDs have a preferential interaction with water, and thus, a higher hydrophilicity.

Ageing of condensates that start as single-component liquid-like systems and transition into multiphase architectures can be promoted not only by disorder-to-order structural transitions, but also by other changes that strengthen the protein–protein interactions over time; e.g. in the microenvironment (e.g. pH, salt, pressure), in the patterns of post-translational modifications, or in the condensate composition. Regardless, the interplay between the timescales of protein self-diffusion and the accumulation of disorder-to-order transitions (or the key factor strengthening biomolecular interactions) determines the properties of the aged condensate, and can give rise to a wide-range of steady-state morphologies and architectures, beyond those considered here. Gradual maturation of solid-like condensates from liquid-like droplets due to disorder-to-order transitions or protein aggregation, as those reported in Refs. [11, 15–23], requires that the timescales of protein self-diffusion are the fastest. That is, when protein diffusion is slower (or comparable) to the rates at which disorder-to-order transitions occur, we expect to observe the formation a kinetically-arrested nuclei that grows into an aspherical solid-like structure. Ageing of liquid condensates into the liquid-core/gel-shell or gel-core/liquid-shell architectures we report here, also necessitates that the rate of accumulation of stronger protein–protein interactions is slower than the timescales of protein self-diffusion. Importantly, relative experimental *β*-sheet transition barriers suggest typical timescales of the order of hundreds of nanoseconds [76–78], protein self-diffusion timescales are of the order of hundreds of milliseconds [79], and fluctuation timescales of high-density protein concentration gradients which lead to gradual rigidification of phase-separated condensates are in the range of minutes [33].

## CONCLUSIONS

In this study, we explore the formation of liquid-core/gel-shell and gel-core/liquid-shell multiphase architectures in single-component protein condensates. Combining atomistic simulations with coarse-grained simulations at two resolutions, we predict that an equilibrium homogeneous single-component protein condensate can age into a nonequilibrium multiphase condensate, where a liquid-like phase is in coexistence with an arrested gel-like phase, due to the advent over time of imbalanced homotypic protein–protein interactions. Strikingly, our simulations propose that such critical imbalanced interactions can emerge intrinsically within single-component protein condensates—i.e., even in the absence of chemical modifications or external stimuli—from a gradual accumulation of inter-protein β-sheets. We further find that the specific ordering of the liquid-like and gel-like phases in the condensate is dictated by the molecular organisation of proteins within each of the two different coexisting phases, because that modulates the properties and interfacial free energies of the various interfaces involved (i.e., condensate-solvent and gel-liquid).

During the ageing of single-component FUS condensates, we find that accumulation of disorder-to-order structural transitions among the PLDs, which give rise to inter-protein β-sheet ladders, can sufficiently enhance the strength of PLD–PLD interactions and drive the transformation of the condensate into a nonequilibrium liquid-core/gel-shell multiphase architecture. Furhtermore, we observe that despite the inner liquid-like core and outer arrested gel-like shell being composed of chemically-identical FUS proteins (i.e., only distinguished by the structure of their PLDs and, hence, strength of PLD–PLD interactions), each phase exhibits strikingly different molecular organisation. That is, the liquid-like low-density phase at the core of the FUS multiphase condensates is structurally homogeneous as it is sustained by weak and transient interactions among FUS proteins that can diffuse freely across the whole phase. By contrast, the molecular organization of the arrested gel-like high-density FUS shell is heterogeneous; i.e., PLD regions form a hydrophobic core due to strengthened PLD– PLD interactions that form kinked cross-β-sheets, and the charge-rich RGG2 and RGG3 domains preferentially expose their positively charged side chains to the solvent. Consistently with the liquid-core/gel-shell architecture, we observe that the FUS gel phase has a more hydriphilic interface, due to its higher surface charge density, than the liquid phase. In contrast, in protein systems where the gel phase has equal or lower hydrophilicity than the liquid, we predict the condensates will form instead a nonequilibrium gel-core/liquid-shell architecture, as that organization increases the enthalpic gain for condensate formation and lowers the surface tension of the overall system.

Importantly, our findings suggest that the formation of disorder-to-order protein structural transitions, hence molecular scale processes, can modulate mesoscale phase behavior of protein condensates and lead to the emergence of nonequilibrium multiphase architectures in single-component protein systems during ageing. This finding is significant as it demonstrates that multiphasic organization can arise not only from multicomponent systems that contain two or more different molecular entities (e.g., two different biomolecules), but also from a single-component system that is driven out of equilibrium by the intrinsic onset of imbalanced inter-molecular forces. While in our example, imbalanced inter-molecular interactions are introduced by distinct FUS structural ensembles (FUS-PLD in a fully disordered *versus* an ordered state), there are several other scenarios from which heterogeneity can arise (e.g., amino-acid sequence mutations, application of mechanical forces [16], and posttranslational modifications [53]). The ability of imbalanced inter-molecular forces to drive single-component mixtures towards complex nonequilibrium architectures has been recently demonstrated for a solution of chiral tetramer model molecules that can transition between two enantiomeric states and form steady-state arrested microphase domains [80]—akin to the nonequilibrium aged multiphase condensates we report here for naturally occurring proteins.

The prediction that nonequilibrium multiphase condensates can emerge from single-component protein systems is interesting from a fundamental point of view, as it highlights how structural disorder-to-order transitions can give rise to significant physicochemical diversity within a condensate without changing the chemical make up of its biomolecular components. We speculate that such transformations in single-component systems may have wide-spread physiological and pathological implications; for example, in the establishment of core–shell structures (e.g., in stress granules) [38], in the formation of multilayered compartments of the FUS-family protein TDP-43 with vacuolated nucleoplasm-filled internal space [42], or other hierarchically organized subcellular condensate structures where molecules may undergo a structural change or experience some other chemical or physical alteration [46].

More broadly, the multiscale modelling strategies developed in this study can be extended to probe the underlying thermodynamic mechanisms and kinetics of other complex equilibrium and nonequilibrium condensate architectures, and contribute to shed light on how molecular-level features influence the properties of compartments in cellular function and dysfunction.

## METHODS

In this work, we develop a multiscale approach to connect fine atomistic features of proteins to the process of condensate ageing. Our method combines descriptions of proteins at three levels of resolution: (1) Atomistic PMF simulations of interacting peptides using two different force fields (a99SB-*disp* [66] and CHARMM36m [67]), (2) sequence-dependent coarse-grained simulations of phase-separated protein condensates [63], and (3) a tailored-made minimal model of dynamically ageing condensates where the inter-molecular forces among proteins are non-conservative. Our MD simulations of condensates are done with the Direct Coexistence Method, which simulates the condensed and diluted phases in the same simulation box separated by an interface. Full details on the atomistic potential of mean force simulations, residue-resolution coarse-grained model and Direct Coexistence simulations, estimation of the number of molecular contacts, minimal coarse-grained model, dynamical algorithm and local order parameter, as well as the simulation details for all resolution models are provided in the SI Appendix

## Acknowledgements

The research leading to these results has received funding from the European Reseach Council (ERC) under the European Union’s Seventh Framework Programme (FP7/2007-2013) through the ERC grant PhysProt (agreement no. 337969) (T.P.J.K.), under the European Union’s Horizon 2020 Framework Programme through the Future and Emerging Technologies (FET) grant NanoPhlow (agreement no. 766972) (T.P.J.K., G.K.), the Marie Sk-lodowskaCurie grant MicroSPARK (agreement no. 841466) (G.K.), and the ERC grant InsideChromatin (agreement no. 803326) (R.C.-G.). A.G. acknowledges funding from the EPRSC (EP/N509620/)and the Winton programme. We further thank the Newman Foundation (T.P.J.K.), the Biotechnology and Biological Sciences Research Council (T.P.J.K.), the Herchel Smith Funds (G.K.), the Wolfson College Junior Research Fellowship (G.K.), the Winton Advanced Research Fellowship (R.C.-G.), the Oppenheimer Research Fellowship (J.R.E.), Emmanuel College Roger Ekins Fellowship (J.R.E), and King’s College Research Fellowship (J.A.J). We also acknowledge funding from the Wellcome Trust Collaborative Award 203249/Z/16/Z (T.P.J.K.). The simulations were performed using resources provided by the Cambridge Tier-2 system operated by the University of Cambridge Research Computing Service (http://www.hpc.cam.ac.uk) funded by EPSRC Tier-2 capital grant EP/P020259/1.

## References

[1] A. A. Hyman, C. A. Weber, and F. Jülicher, Annual Review of Cell and Developmental Biology 30, 39 (2014).

[2] Y. Shin, and C. P. Brangwynne, Science 357 (2017).

[3] S. Banani, H. Lee, A. Hyman, and M. Rosen, Nature reviews. Molecular cell biology 18 (2017).

[4] S. Boeynaems, S. Alberti, N. Fawzi, T. Mittag, M. Polymenidou, F. Rousseau, J. Schymkowitz, J. Shorter, B. Wolozin, L. Van Den Bosch, P. Tompa, and M. Fuxreiter, Trends in Cell Biology 28 (2018).

[5] C. P. Brangwynne, P. Tompa, and R. Pappu, Nature Physics 11, 899 (2015).

[6] E. Gomes, and J. Shorter, Journal of Biological Chemistry 294, 7115 (2019).

[7] E. Martin, A. Holehouse, I. Peran, M. Farag, J. Incicco, A. Bremer, G. Royappa, A. Soranno, R. Pappu, and T. Mittag, Science 367, 694 (2020).

[8] P. Li, S. Banjade, H.-C. Cheng, S. Kim, B. Chen, L. Guo, M. Llaguno, J. Hollingsworth, D. King, S. Banani, P. Russo, Q.-X. Jiang, B. Nixon, and M. Rosen, Nature 483, 336 (2012).

[9] A. G. Larson, D. Elnatan, M. M. Keenen, M. J. Trnka, J. B. Johnston, A. L. Burlingame, D. A. Agard, S. Redding, and G. J. Narlikar, Nature 547, 236 (2017).

[10] J. Wang, J.-M. Choi, A. Holehouse, H. Lee, X. Zhang, M. Jahnel, S. Maharana, R. Lemaitre, A. Pozniakovsky, D. Drechsel, I. Poser, R. Pappu, S. Alberti, and A. Hyman, Cell 174 (2018).

[11] A. Molliex, J. Temirov, J. Lee, M. Coughlin, A. Kanagaraj, H. J. Kim, T. Mittag, and J. P. Taylor, Cell 163, 123 (2015).

[12] T. M. Franzmann, and S. Alberti, Journal of Biological Chemistry 294, 7128 (2018).

[13] N. Lorenzo Gotor, A. Armaos, G. Calloni, M. Torrent, R. Vabulas, N. De Groot, and G. Tartaglia, Nucleic acids research 48 (2020).

[14] Z. March, O. King, and J. Shorter, Brain Research 1647 (2016).

[15] A. Patel, H. Lee, L. Jawerth, S. Maharana, M. Jahnel, M. Hein, S. Stoynov, J. Mahamid, S. Saha, T. Franzmann, A. Pozniakovsky, I. Poser, N. Maghelli, L. Royer, M. Weigert, E. Myers, S. Grill, D. Drechsel, A. Hyman, and S. Alberti, Cell 162, 1066 (2015).

[16] Y. Shen, F. S. Ruggeri, D. Vigolo, A. Kamada, S. Qamar, A. Levin, C. Iserman, S. Alberti, P. George-Hyslop, and T. Knowles, Nature Nanotechnology 15, 1 (2020).

[17] T. Murakami, S. Qamar, J. Q. Lin, G. Schierle, E. Rees, A. Miyashita, A. Costa, R. Dodd, F. Chan, C. Michel, D. Kronenberg-Versteeg, Y. Li, S.-P. Yang, Y. Wakutani, W. Meadows, R. Ferry, L. Dong, G. Tartaglia, G. Favrin, and P. George-Hyslop Neuron (2015).

[18] D. Murray, M. Kato, Y. Lin, K. Thurber, I. Hung, S. Mcknight, and R. Tycko Cell 171 (2017).

[19] M. Hughes, M. Sawaya, D. Boyer, L. Goldschmidt, J. Rodriguez, D. Cascio, L. Chong, T. Gonen, and D. Eisenberg Science 359, 698 (2018).

[20] F. Luo, X. Gui, H. Zhou, Y. Li, X. Liu, M. Zhao, D. Li, X. Li, and C. Liu Nature structural molecular biology 25 (2018).

[21] E. Guenther, P. Ge, H. Trinh, M. Sawaya, D. Cascio, D. Boyer, T. Gonen, Z. Zhou, and D. Eisenberg Nature Structural Molecular Biology 25 (2018).

[22] X. Gui, F. Luo, Y. Li, H. Zhou, Z. Qin, Z. Liu, M. Xie, K. Zhao, B. Dai, W. S. Shin, J. He, L. He, L. Jiang, M. Zhao, B. Sun, X. Li, C. Liu, and D. Li, Nature Communications 10 (2019).

[23] M. Kato, T. Han, S. Xie, K. Shi, X. Du, L. Wu, H. Mirzaei, E. Goldsmith, J. Longgood, J. Pei, N. Grishin, D. Frantz, J. Schneider, S. Chen, L. Li, M. Sawaya, D. Eisenberg, R. Tycko, and S. Mcknight, Cell 149, 753 (2012).

[24] P. George-Hyslop, J. Q. Lin, A. Miyashita, E. Phillips, S. Qamar, S. Randle, and G. Wang, Brain Research 1693 (2018).

[25] S. Alberti, and D. Dormann, Annual Review of Genetics 53 (2019).

[26] S. Alberti, and A. Hyman, Nature Reviews Molecular Cell Biology 22, 1 (2021).

[27] T. Harmon, A. Holehouse, M. Rosen, and R. Pappu, eLife 6, e30294 (2017).

[28] J. Espinosa, J. Joseph, I. Sanchez-Burgos, A. Garaizar, D. Frenkel, and R. Collepardo-Guevara, Proceedings of the National Academy of Sciences 117, 201917569 (2020).

[29] R. Jadrich, and K. Schweizer, The Journal of chemical physics 135, 234902 (2011).

[30] S. Qamar, G. Wang, S. Randle, F. S. Ruggeri, J. Varela, J. Q. Lin, E. Phillips, A. Miyashita, D. Williams, F. Ströhl, W. Meadows, R. Ferry, V. Dardov, G. Tartaglia, L. Farrer, G. Schierle, C. Kaminski, C. Holt, P. Fraser, and P. George-Hyslop, Cell 173, 720 (2018).

[31] A. Bock, A. Murthy, W. S. Tang, N. Jovic, F. Shewmaker, J. Mittal, and N. Fawzi, Protein Science 30 (2021).

[32] Z. Monahan, V. Ryan, A. Janke, K. Burke, S. Rhoads, G. Zerze, R. O’Meally, G. Dignon, A. Conicella, W. Zheng, R. Best, R. Cole, J. Mittal, F. Shewmaker, and N. Fawzi, The EMBO Journal 36, e201696394 (2017).

[33] L. Jawerth, E. Fischer-Friedrich, S. Saha, J. Wang, T. Franzmann, X. Zhang, J. Sachweh, M. Ruer, M. Ijavi, S. Saha, J. Mahamid, A. Hyman, and F. Jülicher, Science 370, 1317 (2020).

[34] C. M. Fare, and J. Shorter, Disease Models & Mechanisms 14, dmm048983 (2021).

[35] M. Feric, N. Vaidya, T. Harmon, D. Mitrea, L. Zhu, T. M. Richardson, R. Kriwacki, R. Pappu, and C. P. Brangwynne, Cell 165 (2016).

[36] C. P. Brangwynne, T. J. Mitchison, and A. A. Hyman, Proceedings of the National Academy of Sciences 108, 4334 (2011).

[37] J. A. West, M. Mito, S. Kurosaka, T. Takumi, C. Tanegashima, T. Chujo, K. Yanaka, R. E. Kingston, T. Hirose, C. Bond, et al., Journal of cell biology 214, 817 (2016).

[38] S. Jain, J. R. Wheeler, R. W. Walters, A. Agrawal, A. Barsic, and R. Parker, Cell 164 (2016).

[39] J. Guilléen-Boixet, A. Kopach, A. S. Holehouse, S. Wittmann, M. Jahnel, R. Schlüßler, K. Kim, I. R. Trussina, J. Wang, D. Mateju, et al., Cell 181, 346 (2020).

[40] H. Yu, S. Lu, K. Gasior, D. Singh, S. Vazquez-Sanchez, O. Tapia, D. Toprani, M. S. Beccari, J. R. Yates, S. Da Cruz, et al., Science 371 (2021).

[41] K. Kistler, T. Trcek, T. Hurd, R. Chen, F.-X. Liang, J. Sall, M. Kato, and R. Lehmann, eLife 7 (2018).

[42] B. Schmidt, and R. Rohatgi, Cell Reports 16 (2016).

[43] I. Alshareedah, M. Moosa, M. Raju, D. Potoyan, and P. Banerjee, Proceedings of the National Academy of Sciences 117, 201922365 (2020).

[44] L. D. Gallego, M. Schneider, C. Mittal, A. Romanauska, R. M. G. Carrillo, T. Schubert, B. F. Pugh, and A. Köhler, Nature 579, 592 (2020).

[45] G. A. Mountain, and C. D. Keating, Biomacromolecules 21, 630 (2020).

[46] T. Lu, and E. Spruijt, Journal of the American Chemical Society 142, 2905 (2020).

[47] S. Boeynaems, A. S. Holehouse, V. Weinhardt, D. Kovacs, J. Van Lindt, C. Larabell, L. Van Den Bosch, R. Das, P. S. Tompa, R. V. Pappu, et al., Proceedings of the National Academy of Sciences 116, 7889 (2019).

[48] T. Kaur, M. Raju, I. Alshareedah, R. Davis, D. Potoyan, and P. Banerjee, Nature Communications 12 (2021).

[49] R. Fisher, and S. Elbaum-Garfinkle, Nature communications 11, 4628 (2020).

[50] I. Sanchez-Burgos, J. A. Joseph, R. Collepardo-Guevara, and J. R. Espinosa, Scientifc Reports 11, 15241 (2021).

[51] F. Dar, and R. V. Pappu, Biophysical Journal 118, 213a (2020).

[52] W. Jacobs, and D. Frenkel, Biophysical Journal 112, 683 (2017).

[53] S. Ranganathan, and E. Shakhnovich, bioRxiv (2021).

[54] A. Garaizar, J. Espinosa, J. Joseph, and R. Collepardo, Scientific Reports 12, 4390 (2022).

[55] S. Ranganathan, and E. I. Shakhnovich, Elife 9, e56159 (2020).

[56] G. Dignon, W. Zheng, and J. Mittal, Current Opinion in Chemical Engineering 23, 92 (2019).

[57] M. Paloni, R. Bailly, L. Ciandrini, and A. Barducci, The journal of physical chemistry. B 124 (2020).

[58] Z. Benayad, S. von Bülow, L. S. Stelzl, and G. Hummer, Journal of Chemical Theory and Computation 17, 525 (2020).

[59] W. Zheng, G. Dignon, N. Jovic, X. Xu, R. Regy, N. Fawzi, Y. Kim, R. Best, and J. Mittal, The Journal of Physical Chemistry B 124 (2020).

[60] G. Krainer, T. Welsh, J. Joseph, J. Espinosa, S. Wittmann, E. Csilléry, A. Sridhar, Z. Toprakcioglu, G. Gudiškytė, M. Czekalska, W. Arter, J. Guillén-Boixet, T. Franzmann, S. Qamar, P. George-Hyslop, A. Hyman, R. Collepardo-Guevara, S. Alberti, and T. Knowles, Nature Communications 12 (2021).

[61] G. Dignon, W. Zheng, Y. Kim, R. Best, and J. Mittal, PLOS Computational Biology 14, e1005941 (2018).

[62] A. Garaizar, and J. R. Espinosa, The Journal of Chemical Physics 155, 125103 (2021).

[63] A. J. Joseph, A. Reinhardt, A. Aguirre, P. Y. Chew, K. O. Russell, J. R. Espinosa, A. Garaizar, and R. Collepardo-Guevara, Nat.Comput,. Sci., in press (2021).

[64] A. Šarić, Y. C. Chebaro, T. P. J. Knowles and D. Frenkel, Proceedings of the National Academy of Sciences 111, 17869 (2014).

[65] S. Samantray, F. Yin, B. Kav, and B. Strodel, Journal of chemical information and modeling 60, 6462 (2020).

[66] P. Robustelli, S. Piana, and D. E. Shaw, Proceedings of the National Academy of Sciences 115, E4758 (2018).

[67] J. Huang, S. Rauscher, G. Nawrocki, T. Ran, M. Feig, B. L. De Groot, H. Grubmüller, and A. D. MacKerell, Nature methods 14, 71 (2017).

[68] L. H. Kapcha, and P. J. Rossky, Journal of molecular biology 426, 484 (2014).

[69] T. C. Michaels, A. Šarić, J. Habchi, S. Chia, G. Meisl, M. Vendruscolo, C. M. Dobson, and T. P. Knowles, Annual review of physical chemistry 69, 273 (2018).

[70] A. Ladd, and L. Woodcock, Chemical Physics Letters 51, 155 (1977).

[71] R. Fernández, J. Abascal, and C. Vega, The Journal of chemical physics 124, 144506 (2006).

[72] J. Espinosa, E. Sanz, C. Valeriani, and C. Vega, The Journal of chemical physics 139, 144502 (2013).

[73] T. J. Welsh, G. Krainer, J. R. Espinosa, J. A. Joseph, A. Sridhar, M. Jahnel, W. E. Arter, K. L. Saar, S. Alberti, R. Collepardo-Guevara, and T. P. Knowles, Nano Letters 22, 612 (2022).

[74] F. Luo, X. Gui, H. Zhou, J. Gu, Y. Li, X. Liu, M. Zhao, D. Li, X. Li, and C. Liu, Nature Structural & Molecular Biology 25, 341 (2018).

[75] E. L. Guenther, Q. Cao, H. Trinh, J. Lu, M. R. Sawaya, D. Cascio, D. R. Boyer, J. A. Rodriguez, M. P. Hughes, and D. S. Eisenberg, Nature Structural & Molecular Biology 25, 463 (2018).

[76] S. V. Kathuria, L. Guo, R. Graceffa, R. Barrea, R. P. Nobrega, C. R. Matthews, T. C. Irving, and O. Bilsel, Biopolymers 95, 550 (2011).

[77] J. J.-T. Huang, R. W. Larsen, and S. I. Chan, Chemical Communications 48, 487 (2012).

[78] J. Kubelka, J. Hofrichter, and W. A. Eaton, Current opinion in structural biology 14, 76 (2004).

[79] A. R. Strom, A. V. Emelyanov, M. Mir, D. V. Fyodorov, X. Darzacq, and G. H. Karpen, Nature 547, 241 (2017).

[80] B. Uralcan, T. J. Longo, M. A. Anisimov, F. H. Stillinger, and P. G. Debenedetti, The Journal of Chemical Physics 155, 204502 (2021).

