## Supplementary material for "Can single-component protein condensates form multiphase architectures?": SI Appendix

### Supplementary Information Appendix: Can single-component protein condensates form multiphase architectures?

Adiran Garaizar[1,*], Jorge R. Espinosa[1,*], Jerelle A. Joseph [1,2,3], Georg Krainer[4], Yi Shen[5,6], Tuomas P. J. Knowles[1,4] and Rosana Collepardo-Guevara[1,2,3]
[1] *Maxwell Centre, Cavendish Laboratory, Department of Physics, University of Cambridge, J J Thomson Avenue, Cambridge CB3 0HE, United Kingdom* [2] *Department of Chemistry, University of Cambridge, Lensfield Road, Cambridge CB2 1EW, United Kingdom* [3] *Department of Genetics, University of Cambridge, Downing Site, Cambridge CB2 3EH, United Kingdom* [4] *Centre for Misfolding Diseases, Yusuf Hamied Department of Chemistry, University of Cambridge, Lensfield Road, Cambridge CB2 1EW, United Kingdom* [5] *School of Chemical and Biomolecular Engineering, The University of Sydney, Sydney, NSW 2006, Australia* [6] *The University of Sydney Nano Institute, The University of Sydney, Sydney, NSW 2006, Australia, *=contributed equally*
(Dated: 25th March 2022)

#### I. POTENTIAL OF MEAN FORCE CALCULATIONS

We estimate the potential of mean force (PMF) by performing a set of atomistic umbrella sampling MD simulations for three different FUS LARKS 4-peptide systems, with sequences: 37-SYSGYS-42, (PDB code: 6BWZ), 54-SYSSYGQS-61 (PDB code: 6BXV) and , 77-STGGYG-82 (PDB code: 6BZP) [1]. The simulations were performed with explicit water and ions using the a99SB-*disp* force field for protein/water/ions[2] using the GROMACS 2018 package [3]. The starting configurations for each simulations consisted of four stacked peptides (two pairs of peptides, stacked on top of each other forming a ladder) taken from the cross-$\beta$-fibril structures resolved crystallographycally [1]. Acetyl and N-methyl capping groups were added to the termini of each peptide. For each 4-peptide system, we performed two sets of simulations. First, we computed the interactions among the four peptides when they remain in the cross-$\beta$-fibril structures by imposing positional restraints on all the heavy atoms (1000 kJ mol$^{-1}$ nm$^{-2}$) in all directions (except for the dissociating peptide in the pulling direction); this ensures that the four individual peptides maintain their secondary structure during the simulation (nevertheless, we have checked that at closer distances than 0.7nm, i.e. before overcoming the dissociation free energy barriers shown in Figure 2B, C and D of the main text; green curves), the 4-peptide structured motif can be stable without positional restraints). On the other hand, for the interactions among disordered peptides, we allow all peptides to freely sample their configurational landscape. In both cases, we kept the positions of the central $C_\alpha$ atoms fixed to control the relative distances between peptides (except in the direction of the reaction coordinate for the dissociating peptide).

As the reaction coordinate, we used the distance between the center-of-mass (COM) of the 'dissociating' peptide and the peptide closest to it in the sampling direction. In the case of the disordered peptides, rather than the COM distance, we took the distance between the central $C_\alpha$'s in the aforementioned chains, to avoid noise due to changes in the COM from fluctuations in the peptide conformations. We spaced windows approximately 0.2Å from one another along the reaction coordinate, from 4 to 27Å (where longest range interactions completely vanish). To sample the steep potential among rigid structures, a spring constant of 24000 kJ mol$^{-1}$ nm$^{-2}$ is used.

For the ordered peptides, we ran simulations of about 30 ns per umbrella window. This guarantees a statistical uncertainty in the binding free energy of about 3.5 $k_B$T (see Figure S2). For the disordered peptides, each window was run as an independent simulation 5 times to ensure adequate sampling. Each independent simulation was run for 60 ns (with a different initial velocity distribution)—i.e., an aggregate simulation time of 300 ns. Such timescale guarantees uncertainties of 1-1.5 $k_B$T at most in the dissociation profile of the disordered LARKS (Figs. 2 (main text) and S1). For the integration of the equations of motion, we used the Verlet algorithm with a time step of 1 fs. The statistical uncertainty of the intrinsically disordered peptide dissociation curves was estimated by computing the standard deviation from 5 independent trajectories, while for the case of the structured peptides, it was computed by applying a bootstrapping method to every single PMF trajectory of the whole dissociation profile [4, 5]. To constrain bond lengths and angles, we used LINCS algorithm [6] with an order of 8 and 2 iterations. The cut-off for the Coulomb and van der Waals interactions were chosen at a conservative value of 1.4 nm. For electrostatics, we used Particle Mesh Ewald (PME)[7] of $4^{th}$ order with a Fourier spacing of 0.12 nm and an Ewald tolerance of $1.5 \cdot 10^{-5}$. The simulations were performed in the $NpT$ ensemble. The temperature and pressure were kept constant using a Nosé–Hoover thermostat [8] at T = 300 K (with 1 ps relaxation time) and a Parrinello–Rahman[9] isotropic barostat at p = 1 bar (with a 1 ps relaxation time), respectively. A NaCl concentration of 0.1 M was used throughout.

Each simulation system comprised a box of approximately 12 x 5.6 x 5.6 nm$^3$. We used ∼12700 water molecules and 24 NaCl ion pairs together with 4 peptides. All our systems were electroneutral (i.e., the total charge was zero). After solvation of the peptides, we performed an energy minimization with a force tolerance of 1000 kJ mol$^{-1}$ nm$^{-1}$, followed by a short, 1000 ps $NpT$ equilibration both with positional restraints (10000 kJ mol$^{-1}$ nm$^{-2}$) for the heavy atoms of the chains in all three directions of space. The analysis of the simulations was carried out using the WHAM [10] analysis implemented in GROMACS [3]. The first 10% of the simulations was discarded as equilibration time; although we note that including it had negligible effects on the PMF results. Following the work of Samantray *et al.* [11], and to confirm that our PMF predictions for the binding interaction strength of ordered *vs.* disordered LARKS motifs are not significantly model dependent, we have repeated our PMF simulations with the CHARMM36m force field [12] (Figure S1) employing the same system sizes and conditions aforementioned for the a99SB-*disp* [2] simulations.

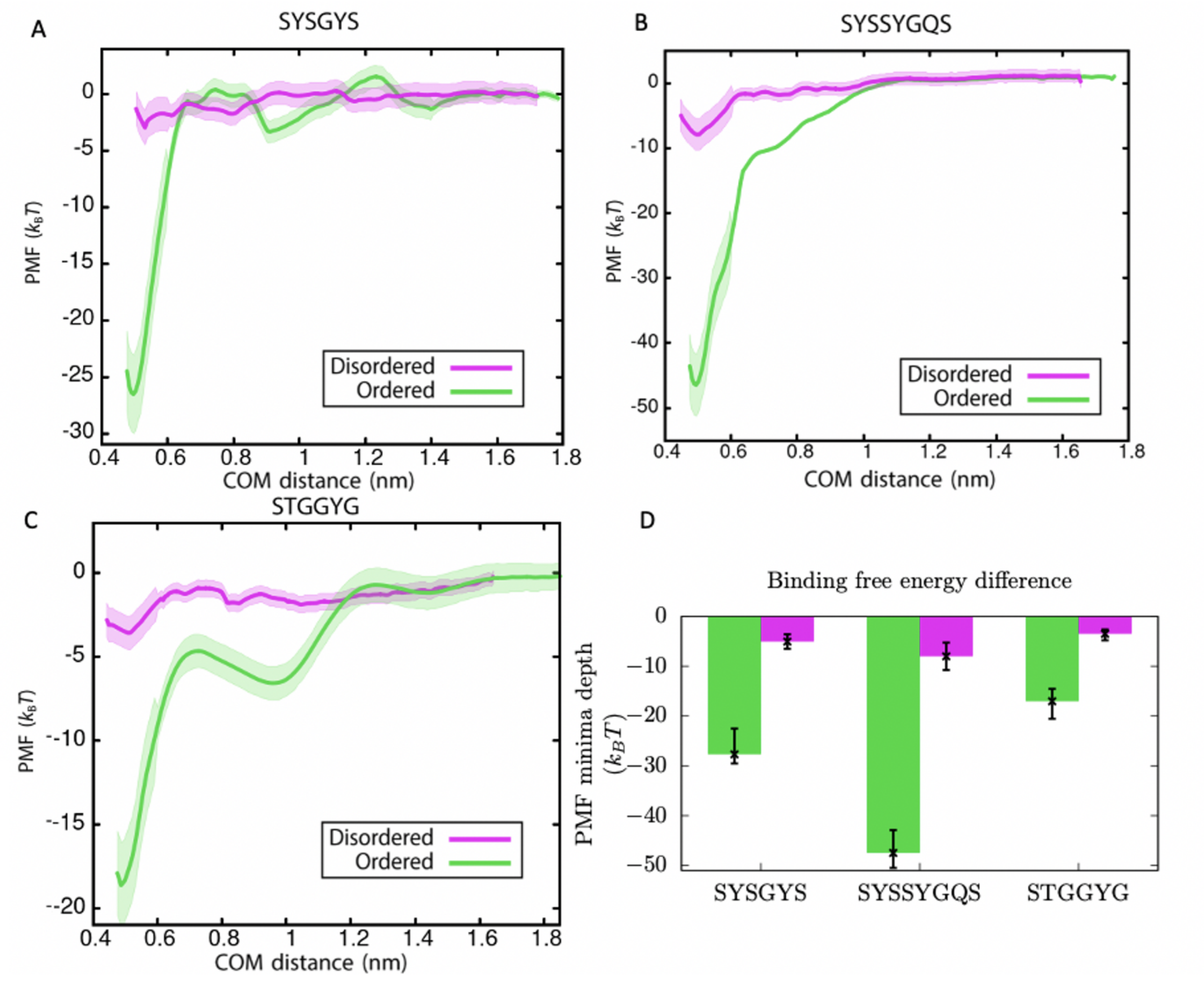


Figure S1: (a–c) Plots of all-atom potential of mean-force (PMF) calculations *versus* centre-of-mass (COM) distance for homotypic pairs of FUS LARKS-forming peptides SYSGYS (PDB code: 6BWZ), SYSSYGQS (PDB code: 6BXV) and STGGYG (PDB code: 6BZP), respectively, before (magenta) and after (green) undergoing the disorder-to-order structural transition. Statistical errors, mean ± standard deviation, are shown as bands; obtained by bootstrapping results from n = 5 independent simulations. The force field employed for these calculations was CHARMM36m [12], and the conditions of the simulations ($T$, $p$ and salt concentration) the same as those shown in Fig. 2 of the main text, which are described in the Materials and Methods Section. (e) Variation in the free energy minimum (as obtained from the profiles shown in panels a–c)).

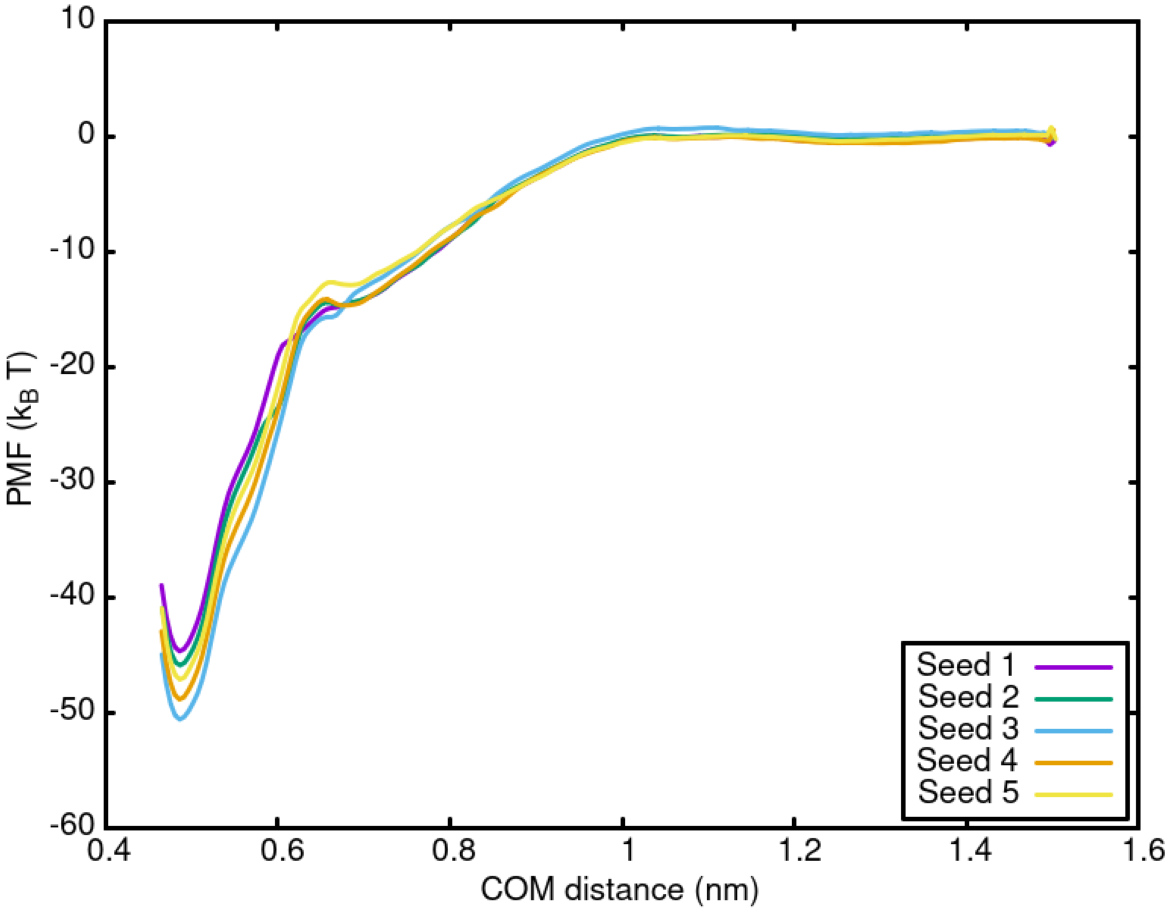


Figure S2: Potential of mean force (PMF) profile for five different independent trajectories as a function of the centre of mass (COM) distance for dissociating a structured peptide from a 4-stacked cross-$\beta$-sheet motif. The sequence of the four peptides was SYSSYGQS (PDB code: 6BXV). Each curve has been computed from a 30-ns independent trajectory once every configuration at every distance interval was fully equilibrated. The conditions ($T$, $p$ and NaCl concentration) of these simulations correspond to those of Fig. 2 of the main text described in the Materials and Methods section. The simulations were performed with the a99SB-*disp* force field [13].

#### II. RESIDUE-RESOLUTION COARSE-GRAINED SIMULATIONS

For our residue-resolution simulations of FUS proteins, we employ the residue-resolution sequence-dependent coarse-grained model Mpipi [14], which describes almost quantitatively the temperature-dependent phase behaviour of different protein solutions. In addition, this model correctly predicts the multiphase behaviour of the PolyR/PolyK/PolyU system, and recapitulates experimental phase separation trends for sequence mutations on FUS, DDX4 NTD and LAF-1 RRG domain variants [14]. Within this force field, electrostatic interactions are modelled with a Coulomb term with Debye–Huckel electrostatic screening [15], while hydrophobic, cation-$\pi$ and $\pi$-$\pi$ interactions are modelled via the Wang–Frenkel potential [16]. Further details on the force field parameters and potentials are provided in Ref. [14] and below.

To set up these residue-resolution simulations, an initial atomistic model of the full-length FUS protein (Uniprot code: K7DPS7, 526 residues) was first developed in VMD [17] fusing the IDRs with the atomistic structures of the globular domains of FUS (i.e., PDBs 2LCW and 6G99). The complete atomistic model was then coarse-grained at the residue-level by placing one bead in each of the $C_\alpha$ atom positions of FUS. For the folded protein domains, we develop an atomistic model by fusing the intrinsically disordered regions of full-length FUS (Uniprot code: K7DPS7, 526 residues) with the resolved structural domains (residues from 285–371 (Protein Data Bank (PDB) code: 2LCW) and from 422–453 (PDB code: 6G99)). Because the model distinguishes between disordered and globular protein regions and maintains the secondary structure of the globular regions (using a rigid body integrator, Ref. [18]), we scale-down by a 30% the set of the Mpipi model parameters (as proposed in Refs. [5, 14]) to account for the buried amino acids belonging to the globular domains. A solution of 48 copies of this coarse-grained FUS proteins was prepared to perform Direct Coexistence simulations [19, 20] at constant volume and temperature. We started by running a simulation of the condensate in the $NpT$ ensemble. After equilibration, we extended one of the sides of the box and then equilibrated in the $NVT$ ensemble. The simulation temperatures were chosen to be slightly below the critical temperature ($T_c$) $T \sim 0.9T_c$ of the model ($T_c = 365\,K$). The production runs were performed for $2\,\mu s$, using a Nosé–Hoover thermostat [8] with a relaxation time of 5 ps and a time step of 10 fs. The LAMMPS software molecular dynamics package was used to carry out all our coarse-grained simulations [18].

##### A. Residue-resolution coarse-grained force field: Mpipi model

Within this force field, Mpipi [14] model, electrostatic interactions are modelled with a Coulomb term with Debye–Huckel electrostatic screening [15], given by the sum over all particle-particle ($i$,$j$) interactions as:

$$E_{elec} = \sum_{i,j} \frac{q_i q_j}{4\pi\epsilon_r\epsilon_0 r_{ij}} exp(-\kappa r_{ij}) \tag{S1}$$

where $q$ is the charge, $\epsilon_r = 80$ is the relative dielectric constant of water, $\epsilon_0$ is the electric constant, $\kappa^{-1} = 795$ pm is the Debye screening length, and $r_{ij}$ is the distance separating particles $i$ and $j$. For these interactions, a Coulomb cut-off of 3.5 nm is employed. The non-bonded interactions between protein beads are modelled via the Wang–Frenkel potential [16]

$$E_{WF} = \sum_{i,j} \epsilon_{ij}\alpha_{ij} \left[\left(\frac{\sigma_{ij}}{r_{ij}}\right)^{2\mu_{ij}} - 1\right] \left[\left(\frac{R_{ij}}{r_{ij}}\right)^{2\mu_{ij}} - 1\right]^{2\nu_{ij}} \tag{S2}$$

where

$$\alpha_{ij} = 2\nu_{ij} \left(\frac{R_{ij}}{\sigma_{ij}}\right)^{2\mu_{ij}} \left[\frac{2\nu_{ij}+1}{2\nu_{ij}\left[\left(\frac{R_{ij}}{\sigma_{ij}}\right)^{2\mu_{ij}} - 1\right]}\right]^{2\nu_{ij}+1} \tag{S3}$$

representing $\sigma$ the molecular diameter of each residue (or nucleotide) and $\epsilon$ the interaction strength between distinct amino acids and nucleotides ($i$ and $j$). $\mu_{ij}$ and $R_{ij}$ are constant model parameters set to $\mu_{ij} = 1$ and $R_{ij} = 3\sigma_{ij}$ for every interaction, while $\sigma_{ij}$ and $\epsilon_{ij}$ are specified for each pair of interaction in Ref. [14]. Finally, bond energy is computed with an harmonic bond potential of the following form:

$$E_{bond} = \sum_{b} \frac{1}{2} k(r_b - r_0) \tag{S4}$$

where $b$ is the total number of bonds along proteins, $r_b$ is the bond distance, $k$=8.03 Jmol$^{-1}$pm$^{-2}$ is the spring constant, and $r_0$ is the bond reference position, set to 381 pm for protein bonds. All the details on this force field and the full list of the model parameters please are provided in Ref. [14].

#### III. MINIMAL COARSE-GRAINED SIMULATIONS

We developed a minimal model for FUS proteins integrating our findings from both the atomistic (Figure 2; main text) and the residue-resolution simulations (Figure 3; main text). We modelled FUS proteins as 20-bead Lennard–Jones (LJ) polymers with one bead representing ca. 26 amino acids; i.e., 6 beads for FUS-PLD, and 14 beads for the RGG1, RRM, RGG2, ZF, and RGG3 regions. A ratio of 6/20 PLD versus total FUS beads recapitulates the ratio of PLD versus total FUS amino acids (i.e., 163/526). The solvent is explicitly modelled using single-bead LJ particles that mimic water-water and water-ion interactions.

Beads that are not directly bonded to each other, establish non-bonded interactions described via the LJ potential:

| | Solvent | PLD ordered | PLD disordered | Non-PLD ordered | Non-PLD disordered |
|---|---|---|---|---|---|
| Solvent | 3.50 | 2.2 | 1.07 | 1.15 | 1.15 |
| PLD (ordered FUS) | 2.20 (1.15) | 3.8 | 1.20 | 1.50 | 1.20 |
| PLD (disordered FUS) | 1.07 (1.15) | 1.2 | 1.45 | 1.25 | 1.25 |
| Non-PLD (ordered FUS) | 1.15 | 1.50 | 1.25 | 1.50 | 1.25 |
| Non-PLD (disordered FUS) | 1.15 | 1.2 | 1.25 | 1.25 | 1.25 |

Table S1: Model parameters for the different $\epsilon$ values in reduced units. Note that the values in parentheses are those for the control simulations in Figure 6 (main text).

| | Solvent | PLD ordered | PLD disordered | Non-PLD ordered | Non-PLD disordered |
|---|---|---|---|---|---|
| Solvent | 1.00 | 1.15 | 1.15 | 1.15 | 1.15 |
| PLD (ordered FUS) | 1.15 | 1.30 | 1.30 | 1.30 | 1.30 |
| PLD (disordered FUS) | 1.15 | 1.30 | 1.30 | 1.30 | 1.30 |
| Non-PLD (ordered FUS) | 1.15 | 1.30 | 1.30 | 1.30 | 1.30 |
| Non-PLD (disordered FUS) | 1.15 | 1.30 | 1.30 | 1.30 | 1.30 |

Table S2: Model parameters for the different $\sigma$ values in reduced units.

$$V(r) = 4\epsilon\left[\left(\frac{\sigma}{r}\right)^{12} - \left(\frac{\sigma}{r}\right)^{6}\right]$$

where $\epsilon$ is the depth of the LJ potential, $r$ the distance between two beads, and $\sigma$ the molecular diameter of each bead. The mass of every bead was chosen to be $m^* = 1$. The parameters ($\epsilon$, $\sigma$) for the various cross-interactions are shown in Tables S1 and S2. From here on, we employ reduced units, which are defined as: $T^* = k_{\rm B}T/\epsilon$, $\rho^* = (N/V)\sigma^3$, $p^* = p\sigma^3/\epsilon$, and time as $\sigma\sqrt{(m/\epsilon)}$.

The cut-off of the LJ interactions is set to 3 times the value of $\sigma$. Bonds between consecutive beads are modelled with a harmonic potential $V_{\rm har} = K(r - r_0)^2$ of $K = 40\ \epsilon/\sigma^2$, and a resting position of $r_0 = 1.3\sigma$. Furthermore, we apply an angular potential of the form, $V_{\rm ang} = K_\theta\ (\theta–\theta_0)^2$, between consecutive bonds, with an angular constant of $K_\theta = 0.2\ \epsilon/rad^2$ and a resting angle of $\theta_0 = 180°$ for fully disordered FUS replicas, and with a constant of $K_\theta = 4\ \epsilon/rad^2$ and $\theta_0 = 180°$ for FUS with ordered PLD, to account for the partial rigidification of the proteins after exhibiting the disorder-to-order fibrillar transition. The model has been parameterized to recapitulate the observed behaviour in FUS proteins with both disordered-like and structured-like PLD interactions (Figs. 3 (main text) and S7) and to reproduce the relative density between water and FUS condensates [5, 21, 22].

Typical system sizes in the minimal model simulations contained 12300 solvent particles and range from 570 to 1088 protein replicas. We used a mixture of 50 mol% fully disordered FUS proteins and 50 mol% with ordered PLD replicas. $NVT$ simulations were run at $T^* = 3.5$ and at density of $p^* = 0.25$. Temperature was kept constant with a Nosé–Hoover thermostat [8] and with a relaxation time of $t^* = 0.4$. The Verlet equations of motion are integrated with a timestep of $t^* = 0.0004$. We employ the Direct Coexistence method [19, 20] as described in the previous section and the LAMMPS Molecular Dynamics software [18].

##### A. Dynamic minimal coarse-grained model

To dynamically mimic [23] the structural diversity of FUS proteins during ageing, we developed a time-dependent minimal coarse-grained algorithm. Based on the above minimal coarse-gained parametrization, we started with a system composed of a homogeneous single phase of FUS proteins with fully disordered PLDs. Then, FUS-PLD domains can spontaneously transition from their fully disordered state to the ordered state depending on the local environment. Every 100 simulation timesteps, the algorithm evaluates whether the conditions around each fully disordered FUS protein are favorable for undergoing an ‘effective’ disorder-to-order cross-$\beta$-fibrillar transition (the exact conditions are described below), and thus modify the protein parameters given in Tables S1 and S2, to those corresponding to ordered PLD-FUS proteins. These conditions are evaluated on the central bead of the PLD, which is a good proxy for the average position of the LARKS found in the PLD of FUS.

The algorithm changes the identity of two FUS chains from the fully disordered state to the ordered PLD interaction parameters once the two following conditions are met: (1) Their two central beads are at a smaller distance than $2.75\,\sigma$, and (2) both central beads are surrounded, within a cut-off distance of $2.75\,\sigma$ (a sensible distance close to the maximum distance at which FUS-PLD beads can still attractively interact; the potential cut-off is $3.25\,\sigma$, which is also used as a local order parameter cut-off in Figure S4), by at least four other central beads and six solvent particles (i.e., characteristic crowded environment of a FUS-rich liquid phase described in Ref. [24] through all-atom and residue-resolution simulations). Due to the strong interaction strength observed in our PMF atomistic simulations between structured FUS domains (Figure 2; main text), we set the transition towards ordered-PLD FUS replicas to be irreversible. To carry out these simulations, we used the USER-REACTION [25] package of LAMMPS which allowed us to change the topology of the underlying system components on-the-fly. The rest of the simulation details are the same as those described in the previous section.

The criterion that at least four peptides should be in close contact to trigger a disorder-to-order transition was chosen based on the following arguments. Four interacting peptides is the minimal system where all the different types of stacking and hydrogen bonding interactions that stabilize the $\beta$-sheet fibrillar ladder are fulfilled; i.e., two interacting steps of the ladder each made of a pair of $\beta$-sheet peptides. Thus, considering fewer interacting peptides (e.g. only three or two) would severely underestimate the strength of interactions among ordered LARKS in our atomistic simulations, and subsequently, erroneously preclude the formation of kinetically arrested states at the coarse-grained level. Our atomistic simulations show that the strength of interactions among four ordered peptides is already high enough for the peptides to remain stably bound upon thermal fluctuations. Hence, if we made the criterion even more stringent (i.e., by requiring clustering of five or more peptides), the strength of interactions among the system would remain consistent with kinetic arrest at the coarse-grained level. However, an stringent criterion would now render the coarse-grained simulations prohibitive expensive. That is, much longer simulation timescales would be needed to capture the rarer higher-density fluctuations that could result in the spontaneous formation of clusters of five or more peptides (as opposed to the more frequent fluctuations that yield clusters of four peptides).

##### B. Estimation of contact frequencies from residue-resolution coarse-grained simulations

The average number of protein domain contacts within phase-separated condensates were calculated using the MDAnalysis Python library [26]. Amino acids of different protein replicas were considered to be in contact when they were found within a cut-off distance of 0.65 nm (although our conclusions barely depend on the choice of cut-off distance, as long as it is reasonable), and we averaged these contact numbers along the production time of each simulation. The contacts between two given domains were obtained by summing all amino-acid contacts found in those domains. Then, from the average number of contacts between different regions along the simulation, we calculated the fraction of total contacts per domain by normalization with the total number of contacts.

[1] M. Hughes, M. Sawaya, D. Boyer, L. Goldschmidt, J. Rodriguez, D. Cascio, L. Chong, T. Gonen, and D. Eisenberg, Science **359**, 698 (2018).
[2] P. Robustelli, S. Piana, and D. E. Shaw, Proceedings of the National Academy of Sciences **115**, E4758 (2018).
[3] H. Berendsen, D. van der Spoel, and R. Drunen, Computer Physics Communications **91**, 43 (1995).
[4] B. Efron and R. Tibshirani, Statistical Science **1** (1986).
[5] G. Krainer, T. Welsh, J. Joseph, J. Espinosa, S. Wittmann, E. Csilléry, A. Sridhar, Z. Toprakcioglu, G. Gudiškytė, M. Czekalska, W. Arter, J. Guillén-Boixet, T. Franzmann, S. Qamar, P. George-Hyslop, A. Hyman, R. Collepardo-Guevara, S. Alberti, and T. Knowles, Nature Communications **12** (2021).
[6] B. Hess, Journal of Chemical Theory and Computation **4**, 116 (2008).
[7] T. Darden, D. York, and L. Pedersen, Journal of Computational Chemistry **18**, 1463 (1993).
[8] S. Nosé, The Journal of Chemical Physics **81**, 511 (1984).
[9] M. Parrinello and A. Rahman, Journal of Applied physics **52**, 7182 (1981).
[10] J. Hub, B. de Groot, and D. van der Spoel, Journal of Chemical Theory and Computation **6**, 3713–3720 (2010).
[11] S. Samantray, F. Yin, B. Kav, and B. Strodel, Journal of chemical information and modeling **60**, 6462 (2020).
[12] J. Huang, S. Rauscher, G. Nawrocki, T. Ran, M. Feig, B. L. De Groot, H. Grubmüller, and A. D. MacKerell, Nature methods **14**, 71 (2017).
[13] P. Robustelli, S. Piana, and D. E. Shaw, Proceedings of the National Academy of Sciences **115**, E4758 (2018).
[14] A. J. Joseph, A. Reinhardt, A. Aguirre, P. Y. Chew, K. O. Russell, J. R. Espinosa, A. Garaizar, and R. Collepardo-Guevara, Nat. Comput. Sci. , in press (2021).

[15] P. Debye, Physikalische Zeitschrift **24**, 185 (1923).
[16] X. Wang, S. Ramírez-Hinestrosa, J. Dobnikar, and D. Frenkel, Physical Chemistry Chemical Physics **22**, 10624 (2020).
[17] W. Humphrey, A. Dalke, and K. Schulten, Journal of molecular graphics **14**, 33 (1996).
[18] S. Plimpton, J. Comp. Phys. **117**, 1 (1995).
[19] A. Ladd and L. Woodcock, Chemical Physics Letters **51**, 155 (1977).
[20] R. García Fernández, J. L. F. Abascal, and C. Vega, The Journal of Chemical Physics **124**, 144506 (2006).
[21] A. Garaizar and J. R. Espinosa, The Journal of Chemical Physics **155**, 125103 (2021).
[22] A. Bock, A. Murthy, W. S. Tang, N. Jovic, F. Shewmaker, J. Mittal, and N. Fawzi, Protein Science **30** (2021).
[23] G. A. Mountain and C. D. Keating, Biomacromolecules **21**, 630 (2020).
[24] T. J. Welsh, G. Krainer, J. R. Espinosa, J. A. Joseph, A. Sridhar, M. Jahnel, W. E. Arter, K. L. Saar, S. Alberti, R. Collepardo-Guevara, and T. P. Knowles, Nano Letters **22**, 612 (2022).
[25] J. Gissinger, B. Jensen, and K. Wise, Polymer **128** (2017).
[26] R. T. McGibbon, K. A. Beauchamp, M. P. Harrigan, C. Klein, J. M. Swails, C. X. Hernández, C. R. Schwantes, L.-P. Wang, T. J. Lane, and V. S. Pande, Biophysical journal **109**, 1528 (2015).

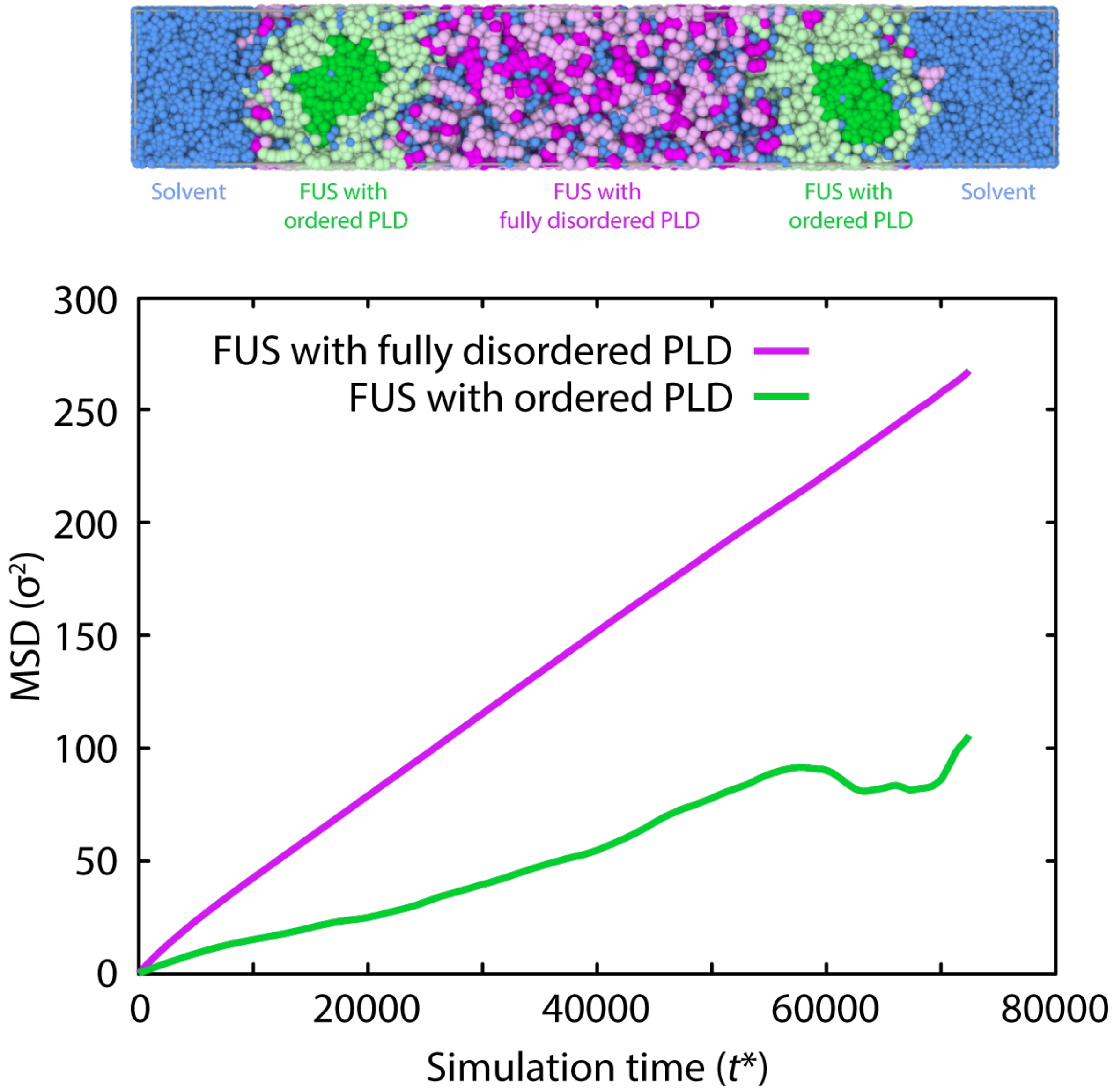


Figure S3: Disorder-to-order transitions in FUS-PLD significantly reduce the diffusion of FUS proteins in our tailored minimal model. Top: Simulation snapshot of the system including FUS proteins with ordered PLD (green chains), FUS proteins with fully-disordered PLD (purple chains), and solvent molecules (blue beads). Please note that darker beads in FUS replicas correspond to the PLD region, while lighter ones to the rest of the protein sequence. Bottom: Mean Squared Displacement *versus* simulation time for FUS proteins with disordered PLDs (purple curve) and for FUS proteins with ordered PLDs (green curve) obtained with our minimal protein model with explicit solvent shown above (see also main text; Fig. 4). As it can be seen, disorder-to-order transitions in FUS-PLD regions greatly hinders protein mobility within the condensates. The mean squared displacement has been evaluated for the central protein bead of each FUS replica within the protein condensate.

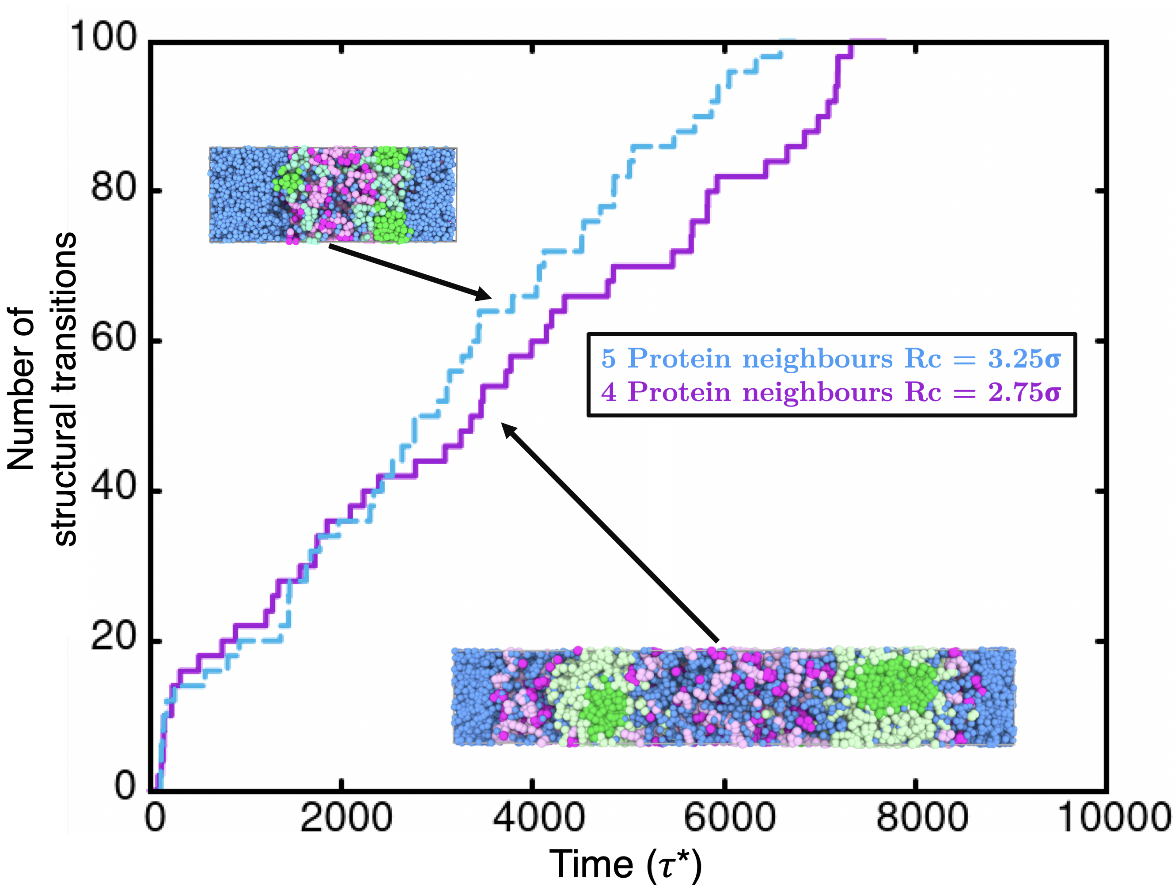


Figure S4: Number of structural transitions as a function of time in the dynamical minimal FUS model with two different local-environment criteria for modulating the emergence of structural transitions in the system. In purple, the original cut-off ($R_c$=2.75$\sigma$) and number (4) of protein PLD neighbours within such cut-off distance employed in the main text simulations to undergo long-lived binding due to the accumulation of inter-protein $\beta$-sheet structures; and in blue, a different cut-off ($R_c$=3.25$\sigma$) distance coupled to a higher number (5) of protein PLD neighbours required to undergo the effective structural transition in our miminal CG model. As it can be seen, an equivalent condensate organization is achieved almost independently of the chosen cut-off distance and the number of protein PLD neighbours as long as that represents a high-local density fluctuation of the protein concentration within the condensate. Representative snapshots of FUS condensates, as obtained via Direct Coexistence MD simulations, are shown. The same colour code described in the caption of Fig. 4 (main text) applies here. Please note that the temperature and pressure of these simulations correspond to those employed for the calculations shown in Figs. 4 and 5 in the main text (and described in the Materials and Methods Section).

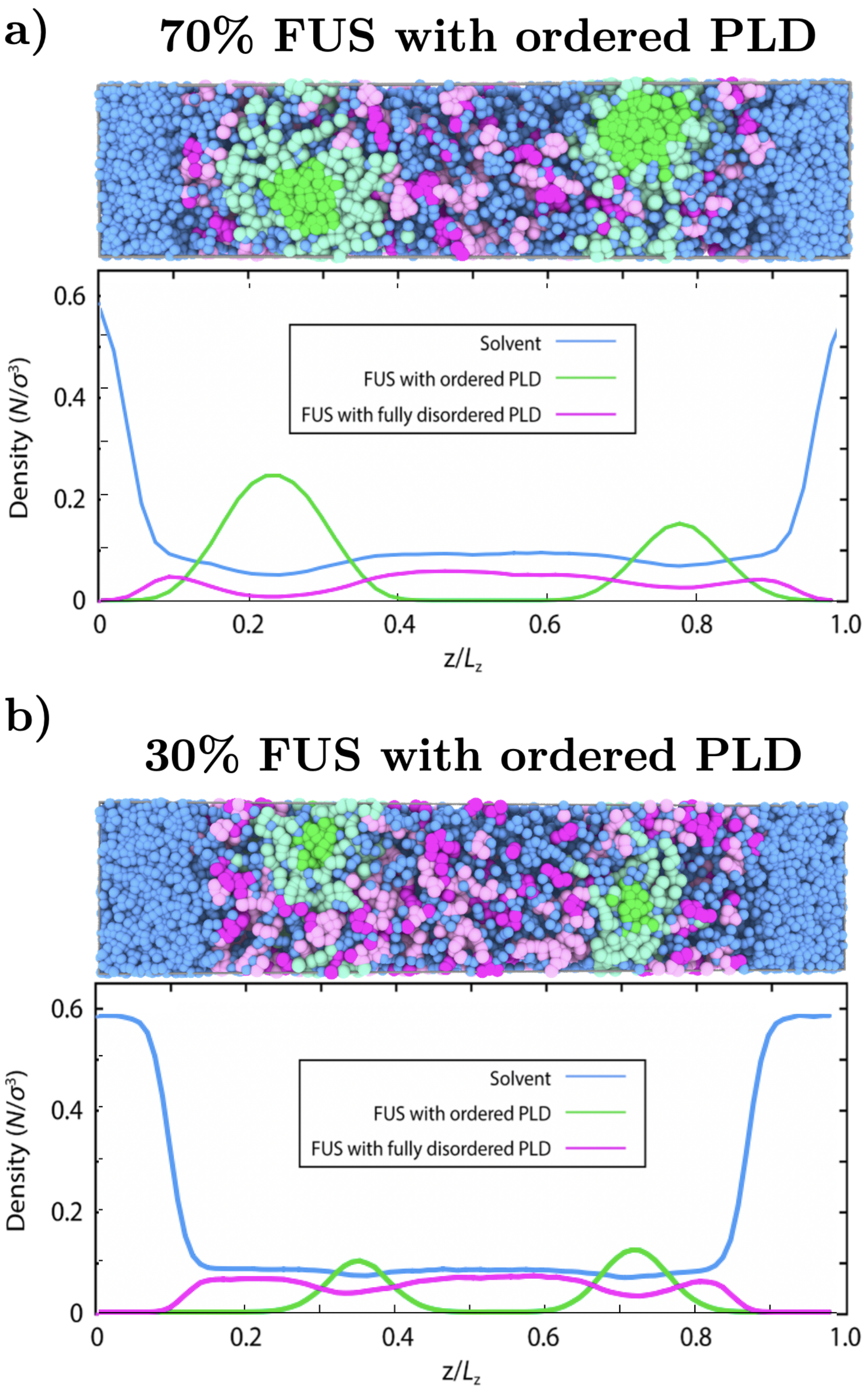


Figure S5: a) Top: Snapshot of the final state of a Direct Coexistence simulation of a mixture of 30% FUS protein replicas with fully disordered PLDs and 70% FUS proteins with cross-$\beta$-sheet elements in their PLDs and explicit solvent. Bottom: Density profile (in reduced units) of FUS species and explicit solvent across the long side of the simulation box estimated over the coarse-grained equilibrium ensemble (as obtained in panel (e) of Fig. 4 in the main text). FUS proteins with fully disordered PLDs: magenta; FUS proteins with ordered PLDs (i.e., with kinked cross--sheets): green, and solvent (water): blue. b) Top: Snapshot of the final state of a Direct Coexistence simulation of a mixture of 70% FUS proteins with fully disordered PLDs and 30% FUS proteins with cross-$\beta$-sheet elements in their PLDs and explicit solvent. Bottom: Density profile (in reduced units) of FUS species and explicit solvent across the long side of the simulation box estimated over the coarse-grained equilibrium ensemble (as obtained in panel (e) of Fig. 4 in the main text and panel (a) of this Figure). The same colour code of panel (a) applies for the Direct Coexistence simulation snapshot and the system density profile shown in panel (b). In total, 1088 FUS protein replicas were included in the simulation. Please note that the temperature and pressure of these simulations correspond to those employed for the calculations shown in Figs. 4 and 5 in the main text (described in the Materials and Methods Section).

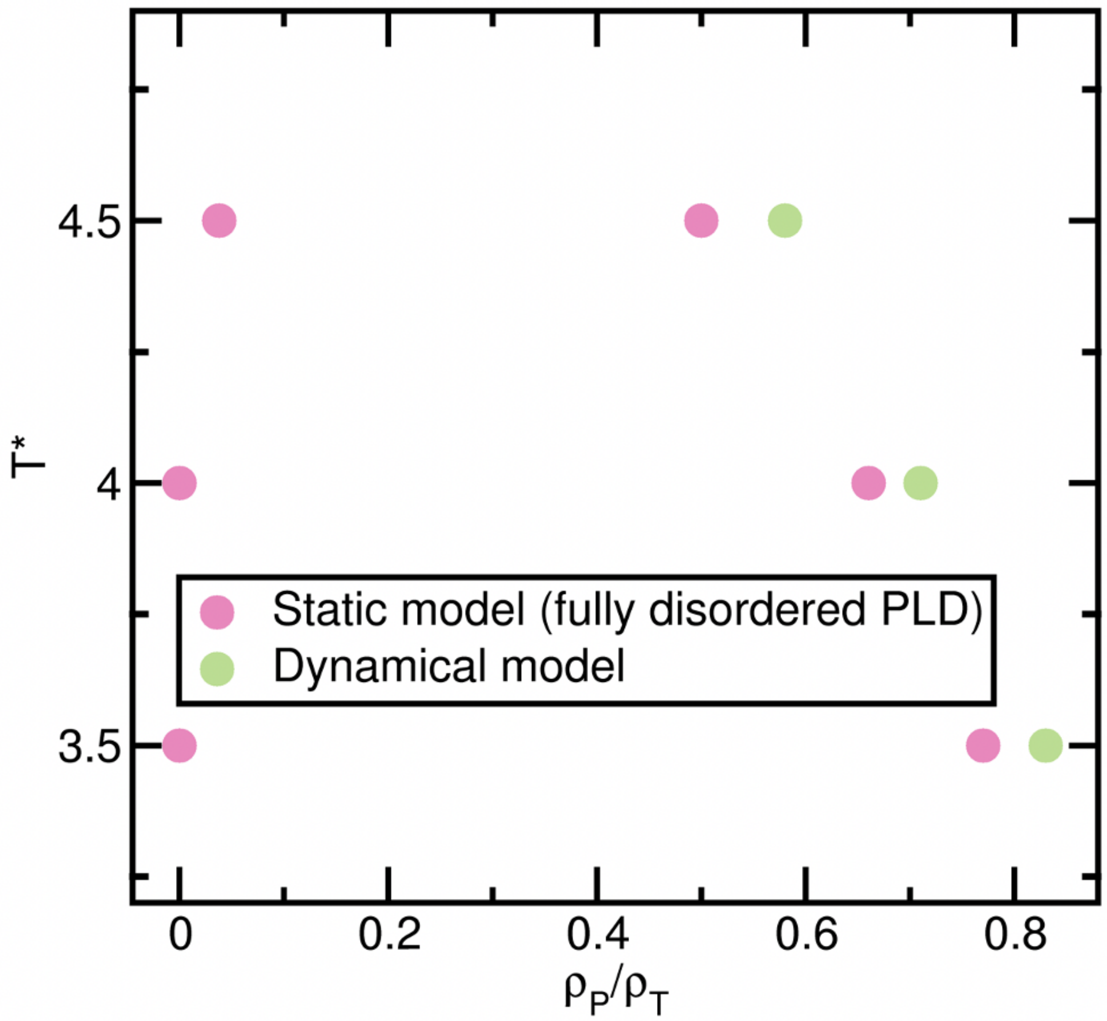


Figure S6: Phase diagram in the $T^*$-($\rho_P/\rho_T$) plane for recently formed FUS condensates in which disorder-to-order structural transitions have not taken place yet (red circles; all FUS-PLDs are set to be disordered), and for aged condensates (in which we switch on the dynamical algorithm and dense gel-like regions emerge on the droplet interfaces while liquid-like low-dense regions remain in the condensate core, as shown in Fig. 5 of the main text). The protein density ($\rho_P$) within the condensate (globally considered; right branch) and the protein density in the dilute phase (left branch) is normalized by the system total density ($\rho_T$). The time window at which the densities were computed for the dynamical model, corresponds to that in which the number of disorder-to-order structural transitions starts to plateau (see inset of Panel (c) in Fig. 5 of the main text).

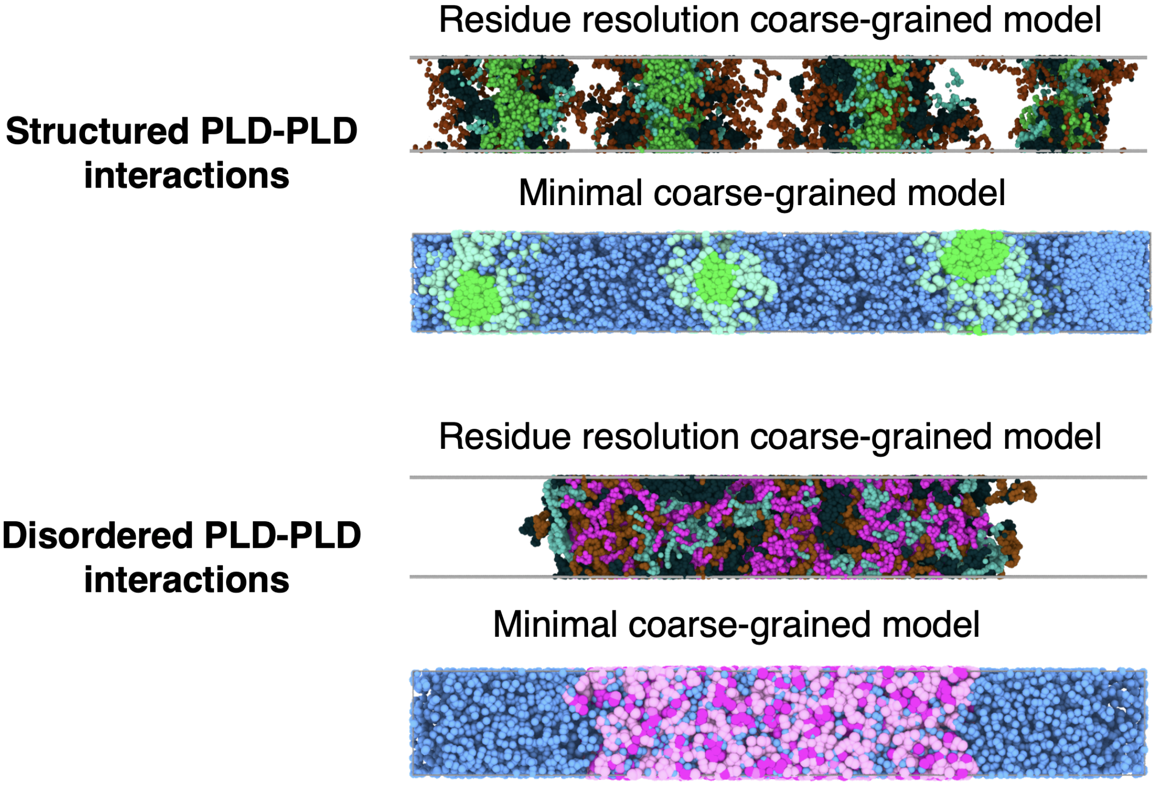


Figure S7: Representative snapshots of FUS replicas, as obtained via Direct Coexistence MD simulations using the Mpipi model [14] and our tailor-made minimal CG model. Amino acid beads in the residue-resolution coarse-grained model are colored according to the domains of FUS, with one bead representing each amino acid: PLD (residues 1–165): green (for structured PLDs) or magenta (for disordered PLDs) ; RGG1 (residues 166–284): cyan; RGG2 (residues 372–422) and RGG3 (residues 454–526): ochre, RRM (residues 285–371) and ZF (residues 423–453): dark blue. In the minimal model simulations, PLD: Magenta (for disordered PLDs) or green (for structured PLDs); RGG1, RRM, RGG2, ZF, RGG3: light magenta or light green according to the type of PLD in the corresponding FUS sequence. Solvent (water) is depicted as blue beads. In the two upper panels, Direct Coexistence simulations possess the reference interaction strength among PLDs (mimicking disordered PLD-PLD interactions) while in the two lower panels the interaction strength among PLDs has been enhanced according to the PMF results shown in Fig. 2e of the main text (mimicking structured PLD-PLD interactions). 48 FUS proteins were included in the Mpipi simulations and 580 FUS replicas in the minimal CG simulations. To obtain the same condensate behavior within the two model resolutions, the two shortest sides (box section; $L_x$ and $L_y$) of the simulation box in the minimal FUS calculations need to be commensurable with the protein size / box dimension ratio used in the residue-resolution simulations (i.e., the shortest side of the simulation box needs to be only between 3-6 times longer than the protein radius of gyration ($R_g$), as required by the computational limitations of residue-resolution simulations). $L_x$=$L_y$=150 Å; FUS $R_g \sim 50$ Å in the Mpipi model; and $L_x$=$L_y$=24 $\sigma$; FUS $R_g \sim 4$ $\sigma$ in the minimal CG model.

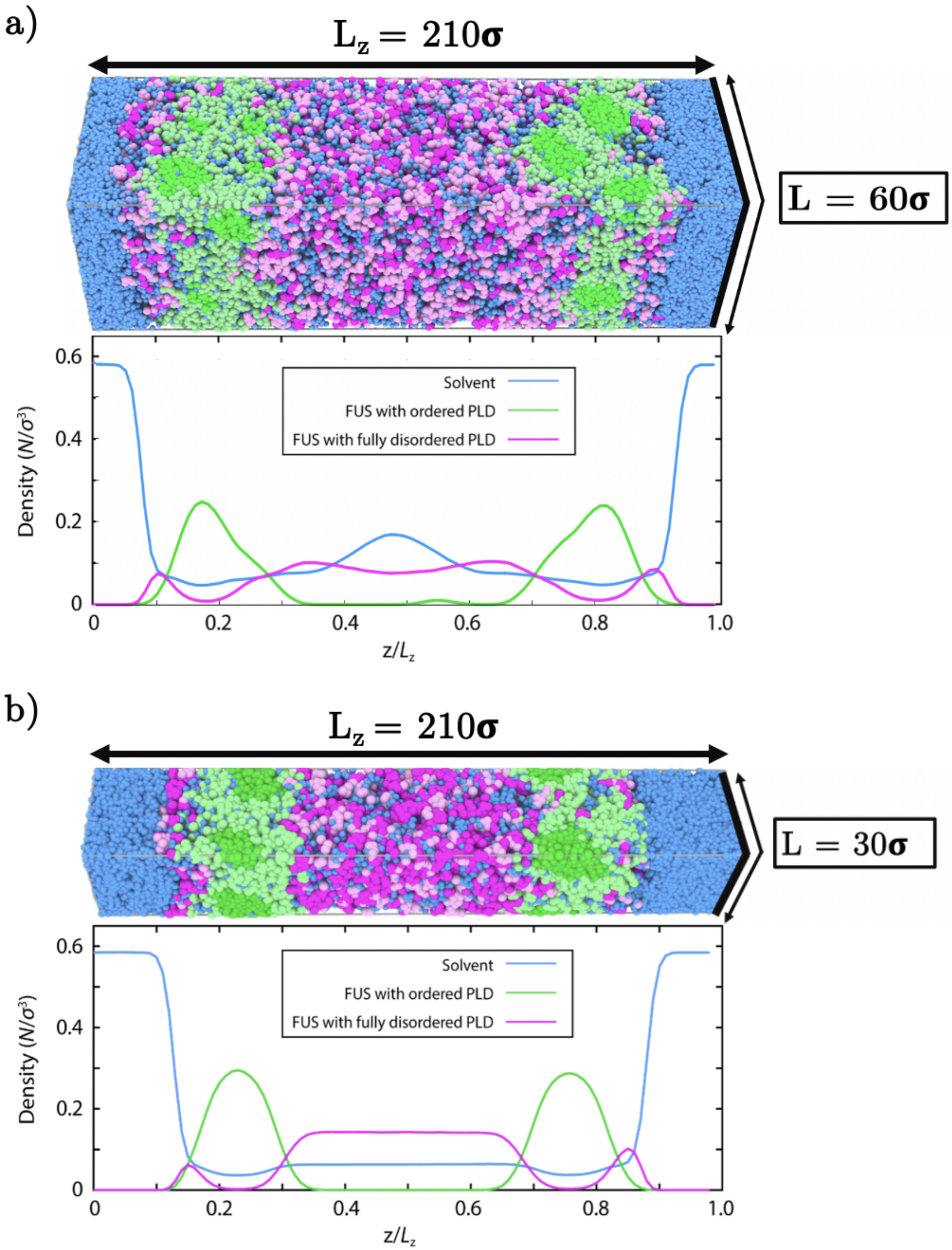


Figure S8: a) Top: Snapshot of a Direct Coexistence simulation using the dynamical algorithm (described in Fig. 5 of the main text) after the emergence of structural transitions within the phase-separated condensate (see inset of Fig. 5 Panel (c)). Bottom: Density profile (in reduced units) of FUS species and explicit solvent across the long side of the simulation box estimated over the coarse-grained multiphase FUS condensate. FUS proteins with fully disordered PLDs: magenta; FUS proteins with ordered PLDs (i.e., with kinked cross-$\beta$-sheet elements): green; solvent (water): blue. The simulation box sides are indicated in the snapshot (please note that $L$=$L_x$=$L_y$). b) The same as in Panel (a) but for a smaller simulation box with a cross-section of $L_x$=$L_y$=30 $\sigma$. Please note that the temperature and pressure of these simulations correspond to those employed for the calculations shown in Figs. 4 and 5 in the main text (and described in the Materials and Methods Section).
